# Refining RNA-Seq Transcriptome Profiles with Transcript-Specific Minimal-Shared Regions

**DOI:** 10.64898/2026.10.28.762701

**Authors:** Qiang Su, Yi Long, Deming Gou, Junmin Quan, Qizhou Lian

## Abstract

Accurate transcript quantification remains a critical challenge in RNA sequencing (RNA-seq) due to ambiguities in read assignment caused by shared regions among transcripts. Beyond shared exons among homologous isoforms, we identified the widespread presence of extensive undefined shared regions-overlapping sequences common to exons or transcripts of heterologous genes-across diverse species, including animals and plants. To overcome this issue, we developed a minimal-sharing strategy that partitions transcripts into unique sets of k-mers and systematically evaluates overlap across all transcripts. By isolating minimal-shared regions specific to each isoform, our approach reduces mapping uncertainty, thereby minimizing false positives and false negatives in transcript detection. Comprehensive evaluations using both in silico simulations and empirical sequencing data demonstrated that our method outperforms traditional probabilistic model-based quantification methods. This strategy improves isoform abundance estimates, enhances gene ontology annotations, and identifies distinct transcript isoforms critical for understanding biological processes, as exemplified in a Huntington’s disease mouse model.

## Background

RNA sequencing (RNA-seq) has transformed our understanding of gene expression and regulation, offering deep insights into the dynamic landscape of transcriptomes.[1, 2] This technique has become essential for precisely investigating a wide range of biological processes by quantifying transcript abundances at the isoform level, aiding in the unraveling of complex cellular mechanisms and disease pathways.[3-6] Recent studies underscore the significance of isoform diversity, which arises from tissue-specific expression and developmental stages, such as those involved in brain development and immune cell differentiation.[7-10] These findings highlight the critical role of alternative splicing in both health and disease, particularly within complex, tissue-specific, and developmental contexts. Currently, RNA-seq predominantly utilizes short-read sequencing due to its high throughput and data quality, enabling the parallelization and error correction necessary for accurately capturing the transcriptome, including low-abundance transcripts.[11-13] However, maximizing the potential of RNA-seq-especially in short-read applications-requires a meticulous workflow.[14, 15] One key challenge in achieving precise isoform-level quantification is the ambiguity in read assignments caused by short sequencing fragments. Since multiple isoforms often share exons, most of reads may align with more than one isoform, leading to unpredictable false positives and false negatives. This imprecision in quantifying individual transcript isoforms can lead to systemic errors in transcriptome abundance quantification.[14, 16] Furthermore, sequencing biases can cause uneven read distribution across transcript template,[17-20] further complicating the accurate quantification of individual isoforms. These issues can significantly mislead downstream biological interpretations.

Despite these challenges, various computational methods have been developed to determine the abundance of individual transcript isoforms from a mixture of sequenced reads. These strategies broadly fall into two categories: alignment-based and alignment-free approaches, both of which rely on optimization algorithms to estimate transcript abundance. Alignment-based methods, such as STAR,[21] or HISAT2,[22] align short RNA-seq reads to a reference genome or transcriptome. In these tools, soft clipping is a standard feature, facilitating the alignment of reads that do not perfectly match the reference sequence. After alignment, reads are assigned to genomic features using tools like Cufflinks,[23] StringTie,[24] and RSEM,[16] producing transcript-level quantification. In contrast, alignment-free methods bypass the alignment step, focusing on sequence composition. For instance, k-mer-based algorithms like Kallisto[25] and Sailfish[26] fragment reads into fixed-length k-mers and count their occurrences. Other tools, such as Salmon,[27] use graph-based representations to analyze k-mers, offering a more flexible and efficient approach to transcript quantification. Both alignment-based and alignment-free approaches fundamentally rely on probabilistic modeling to estimate transcript isoform abundances. Typically, they employ the Expectation-Maximization (EM) algorithm, or its variants, to address the ambiguities in assigning reads or k-mers to specific isoforms.[28, 29] However, the iterative process requires an empirically derived initial estimate of isoform abundances, which can introduce unpredictable biases. Additionally, the uncertainty inherent in probabilistic read assignments poses a persistent challenge, particularly for transcripts with low to moderate expression levels.[23]

In this study, we present a minimal-sharing strategy designed to improve transcriptome quantification by tackling the challenge of mapping uncertainty. This method segments full-length transcript sequences into unique k-mer sets, which are then systematically compared against all possible k-mers. The result is a comprehensive overlap profile spanning the entire genome or transcriptome. Our initial analysis revealed that large segments of reference gene sequences are shared with other, often unrelated, genes. Furthermore, portions of reference transcript sequences significantly overlap with other transcripts. Notably, while overlapping regions are typically thought to stem only from predefined shared exons or homologous transcript isoforms, we found that they also occur extensively across heterologous genes and transcript sequences. Such heterogeneous overlapping inevitably increases mapping ambiguities, leading to large variations in quantification. To address this issue, we screened the transcript-specific overlapping profiles to identify minimal-shared k-mer sets. We evaluated the degree of overlap for each k-mer along the transcript, isolating regions unique to each transcript and excluding areas with complex isoform overlap. Although these minimal-shared regions are shorter than full-length transcripts, they capture the distinct sequence features that define each isoform. Additionally, we refined k-mer counts within these regions using a self-benchmarking system, which helps correct biases and improves the accuracy of transcript quantification. To validate the minimal-sharing strategy, we conducted a comprehensive evaluation of transcriptome overlap profiles across multiple species, including Homo sapiens, Mus musculus, Danio rerio, and the plant Arabidopsis thaliana. Our analysis, based on both in silico simulation data and empirical sequencing data, demonstrated that the minimal-sharing approach consistently outperforms traditional quantification methods. These include alignment-based tools such as Cufflinks, StringTie, and RSEM, as well as alignment-free tools like Salmon and Kallisto, which rely on standard probabilistic models for transcript abundance quantification but can produce unpredictable false positives and false negatives. In an RNA-seq analysis of a Huntington’s disease (HD) mouse model, our minimal-sharing region-based method eliminated these false positives and negatives, improved gene ontology (GO) annotations, and highlighted significant HD-specific isoforms, demonstrating its effectiveness in enhancing transcriptome quantification. Notably, the minimal-sharing strategy seamlessly integrates the identification of differentially expressed isoforms with downstream RNA interference assays, specifically targeting the minimal-shared regions. The complete minimal-sharing code and pipeline for transcriptome quantification based on minimal-shared regions are publicly available on GitHub at https://github.com/QiangSu/Minimal-Sharing-Strategy.

## Results

### A minimal-sharing strategy for resolving ambiguities in transcript quantification

Transcript quantification often faces significant challenges due to ambiguous regions, which complicate the assignment of sequencing reads to specific transcripts. While entire transcript sequences are generally unique, subsegments can share similarities with other transcripts, leading to ambiguity in read assignment. To elucidate this ambiguous region, we employ a k-mer splitting method (as shown in supporting information Fig. S1). By dividing gene exons into sliding k-mer sets (overlapping k-mers with a one-base step) and examining how these k-mers overlap with those from other exons, we can identify shared regions that indicate potential ambiguities. These shared regions suggest possible issues with read assignment during alignment-based quantification (as depicted in Fig. 1a). These ambiguities arise when different, non-related exons (heterologous) share regions, making it difficult to align and count sequencing reads accurately. At the transcriptome level, such k-mer-defined shared regions significantly complicates k-mer matching in alignment-free quantification methods, leading to quantification uncertainties (as illustrated in Fig. 1b). To illustrate these ambiguous genomic and transcriptomic regions more clearly, we firstly highlight a specific shared region on chromosome 7 (positions 5,528,097-5,528,156), corresponding to the ACTB gene in the Homo_sapiens.GRCh38.dna.primary_assembly.fa reference genome. A 50-mer sequence from this exon (ENSE00003465957) is shared by other exons (as shown in Figs. 1c and 1d). Each 50-mer segment within an exon can overlap with segments from other exons (as detailed in Supporting Information Fig. S2). This overlap is quantified by two metrics: global frequency, which measures the extent of overlap across all exons, and local frequency, which assesses overlap within exons defined by a specific gene exon. In the case of exon ENSE00003465957, the shared 50-mer sequences appear not only in other homologous exons of the ACTB gene but also in exons from heterologous genes, such as exon ENSE00002076495 from ACTBP2. Similarly, other exons of the ACTB gene exhibit complex global 50-mer overlap patterns, with sequence uniqueness reflected in a local overlapping degree of 1. Increasing the k-mer length raises the stringency for sequence matching. However, even at a k-mer length of 500, overlaps still occur between 500-mer sequences from one exon and those from other exons (as demonstrated in Supporting Information Fig. S3). These extensive overlaps contribute to substantial ambiguity in sequence assignment during RNA-seq analysis.

**Fig. 1.**
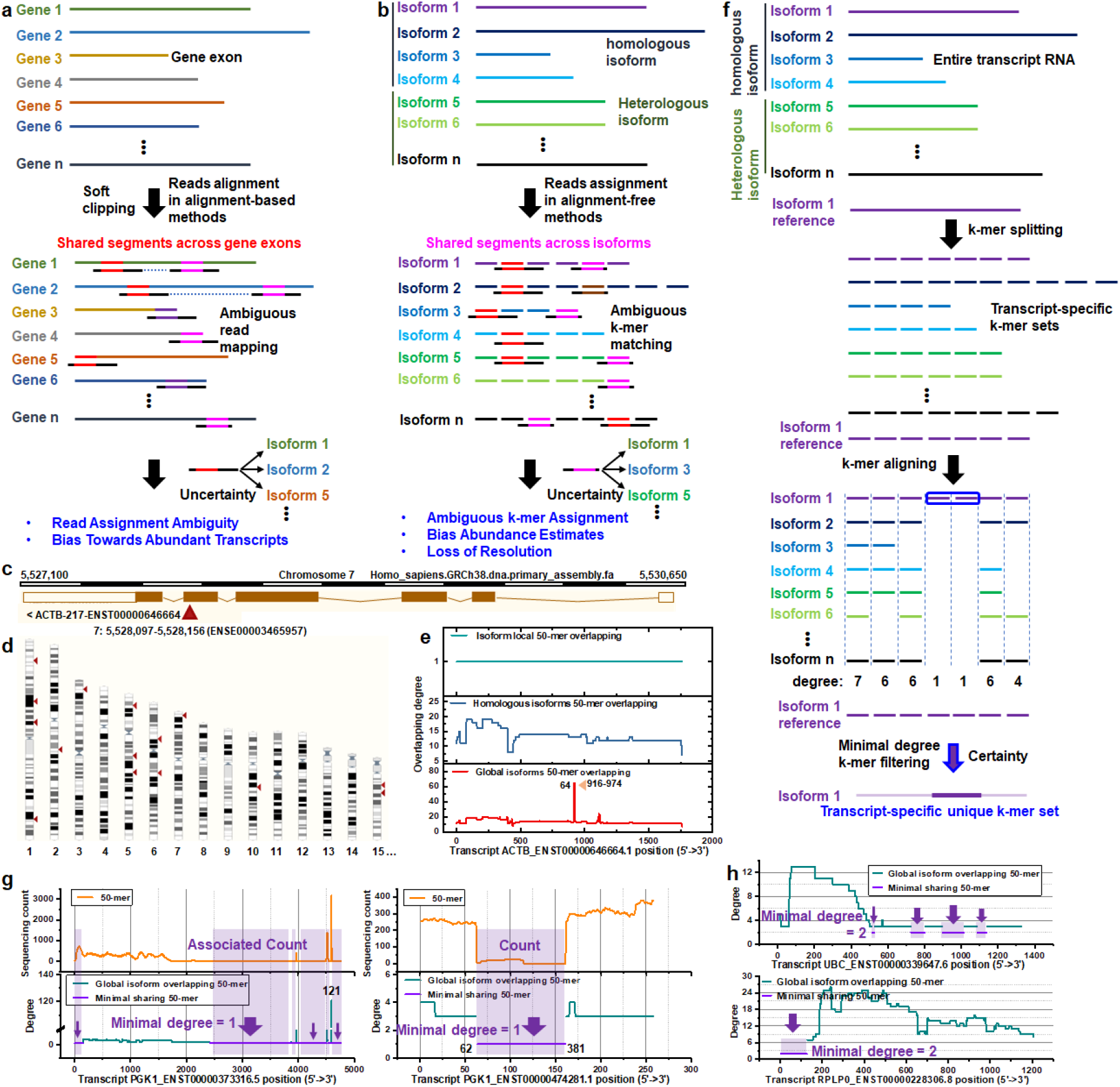
Minimal-sharing strategy for characterizing individual transcripts. **a**) The exon sequence of a gene is segmented into a continuous series of specific k-mers. These gene-specific exon k-mers are then compared with k-mers from other genes to identify shared k-mers, or overlapping regions. In alignment-based transcript quantification methods, the presence of shared sequences across different gene references introduces ambiguity, complicating accurate abundance estimation. **b**) A similar k-mer splitting strategy is applied to the entire transcript isoform sequence, generating isoform-specific k-mer sets. These k-mers are compared with those of other isoforms, and the presence of shared regions complicates k-mer assignments in alignment-free quantification methods. **c-d**) A region within an exon of the ACTB gene (chromosome 7: positions 5,528,097-5,528,155, based on the Homo_sapiens.GRCh38.dna.primary_assembly.fa reference genome) associated with transcript ACTB-ENST00000646664.1 contains a 50-mer that is identical to sequences in other genes located on different chromosomes (Ensemble BLAT search). These shared k-mers exemplify how identical sequences across different genomic locations can result in soft-clipping during sequence alignment. **e**) Each consecutive 50-mer across the full ACTB transcript (ENST00000646664.1) was matched against all other 50-mers from this transcript to assess local overlap. These k-mers were further compared against homologous ACTB isoforms, revealing overlaps, and against all other isoforms, showing global k-mer sharing. A region between positions 916-974 is shared by 63 other isoforms, highlighting the extent of cross-mapping. **f**) The minimal-sharing strategy addresses this ambiguity issue by identifying isoform-specific regions. Each isoform’s k-mers are matched globally against those from other isoforms to generate a k-mer sharing profile. The minimal-shared degree quantifies the uniqueness of the isoform’s sequences. **g**) Two PGK1 isoforms, PGK1_ENST00000373316.5 and PGK1_ENST00000474281.1, are shown with their global 50-mer overlap profiles and corresponding sequencing counts. Regions with minimal sharing (degree = 1) are associated with low sequencing counts, indicating unique isoform-specific regions. **h**) Isoforms such as UBC_ENST00000339647.6 and RPLP0_ENST00000228306.8 lack unique regions, as the minimal-sharing degree excesses 1. A minimal-sharing degree of 2 indicates that each region is shared by a minimum set of two isoforms.

At the transcript level, we analyzed a 50-mer along the ACTB-ENST00000646664.1 reference using three profiles: the global isoform overlapping degree, the homologous isoform overlapping degree, and the local overlapping degree. The global overlapping degree reflects how often a k-mer appears across the entire transcriptome, while the homologous profile shows k-mer occurrences at the gene family level, and the local profile captures k-mers specific to a given isoform. Notably, all 50-mers exhibit highly unique characteristics within the local profile along ACTB-ENST00000646664.1 (as outlined in Fig. 1e). In contrast, the global overlapping profile highlights positioned-50-mers with a higher degree of overlap, with a maximum degree of 64, meaning that the k-mer at this position is shared by 64 different isoforms. This observation suggests that certain transcript segments are shared not only among related homologous isoforms but also with unrelated heterologous genes or transcripts, further complicating ambiguous read assignment. Our comprehensive analysis reveals a consistent pattern across nearly all transcripts, characterized by three distinct features: high uniqueness in local overlapping patterns, moderate complexity in homologous isoform overlapping patterns, and intricate global overlapping patterns (as depicted in Supporting Information Fig. S4). The stark contrast between the global overlapping profile and the homologous overlapping profile underscores the extreme complexity of common-shared regions prevalent in transcript sequences.

To address the challenge of complex isoform overlap, we developed a minimal-sharing strategy to characterize transcript isoforms by focusing on regions with the least overlap (see Fig. 1f). This strategy uses a k-mer splitting method to generate an isoform-specific k-mer set, where each k-mer is matched against those from all other transcripts to create the global overlapping profile. Minimal-shared k-mers are defined as the unique regions of an isoform, offering a clearer distinction for isoform quantification. For instance, when comparing two isoforms of the PGK1 gene, PGK1_ENST00000373316.5 and PGK1_ENST00000474281.1, the impact of this minimal-shared filtering strategy becomes evident (as depicted in Fig. 1g). In PGK1_ENST00000373316.5, the region with the highest overlap degree of 121 corresponds to the area with the highest sequencing read counts. In contrast, a 50-mer region with a minimal overlap degree of 1 represents a unique segment of the isoform. Similarly, the region of PGK1_ENST00000474281.1 spanning positions 62 to 381 exhibits an overlap degree of 1, which is significantly lower than its surrounding regions. This pattern is consistently reflected in the sequencing count profile, where a sharp drop in the read counts further supports the observation of a minimal-shared region. A comparable sharp decline in both the global overlap degree and the sequencing count profile is evident in the PGK1_ENST00000476531.1 homologous isoform (see Supporting Information, Fig. S5). This pattern, characterized by minimal shared regions, appears to be a common feature across most isoform k-mer overlap profiles.

To confirm the consistency of this pattern, we conducted a comprehensive analysis of all six annotated PGK1 isoforms, examining their 50-mer global overlapping degrees and corresponding sequencing counts in parallel (as shown in Supporting Information, Fig. S6). Regions with minimal sharing consistently show the lowest overlapping degrees, creating distinct boundaries with their neighboring high-overlap regions. This pattern strongly correlates with 50-mer sequencing coverage, where regions of high overlap correspond to elevated sequencing counts, and areas of low overlap show reduced coverage. Notably, a 50-mer sequencing count of zero in minimal-shared regions indicates the absence of such isoforms, as observed for PGK1 isoforms ENST00000491291.1, ENST00000644362.1, and ENST00000477335.5. To assess all six isoform abundance, various quantification methods were employed, including alignment-based approaches (Cufflinks, StringTie, and RSEM) and alignment-free tools (Salmon and Kallisto), as well as GaussF (detailed in the Methods section: Transcript Quantification with GaussF), a novel method developed in-house that quantifies transcript abundance using transcript-specific minimal-shared regions (see Supporting Information, Fig. S7). GaussF leverages a theoretically grounded, parametric Gaussian framework, featuring a self-benchmarking algorithm that identifies and corrects systemic biases in RNA-seq data. This correction is performed through k-mer analysis informed by GC content, which guides the construction of a Gaussian distribution essential for comprehensive bias rectification. The incorporation of a pseudo-cumulative distribution function (pseudo-CDF) ensures accurate transcript quantification without relying on empirical initial estimates. The abundance profiles of the PGK1 isoforms determined by GaussF (as depicted in Supporting Information, Fig. S8) showed that isoforms ENST00000491291.1, ENST00000644362.1, and ENST00000477335.5, which have minimal-shared regions with zero sequencing counts, were undetectable by GaussF. StringTie produced similar abundance patterns to GaussF but detected ENST00000491291.1 with a low expression level of 0.1 FPKM, though not zero. In contrast, these three isoforms were detected at notable abundance levels by other methods, raising significant questions about the reliability of different transcript quantification approaches. Both alignment-based and alignment-free traditional methods rely on probabilistic modeling, typically employing the Expectation-Maximization (EM) algorithm or its variants to estimate transcript isoform abundances.[28, 29] These approaches face inherent challenges due to ambiguities in mapping reads or k-mers to specific isoforms, which inevitably introduces uncertainty in transcript quantification.

In transcript isoform analysis, complex overlapping patterns can arise where the minimal-sharing degree among transcript-specific regions exceeds 1. For example, the isoforms UBC_ENST00000339647.6 and RPLP0_ENST00000228306.8 (as presented in Fig. 1h) have a minimal-sharing degree of 2, indicating that certain regions are consistently shared by at least two isoforms. These extensive sharing patterns pose a significant challenge because they make it impossible to identify unique regions for individual reference isoforms. Consequently, quantification based on these minimal-shared regions reflects the combined expression of all involved isoforms. While we can identify the minimal set of isoforms contributing to these shared regions, attempting to distribute reads or k-mer counts among individual isoforms introduces substantial uncertainty. To maintain analytical integrity, we treat these quantifications as composite values rather than assigning them to specific isoforms. This conservative approach is crucial for downstream analyses, where accurate data interpretation depends on acknowledging and accounting for the collective nature of these measurements.

### Assessing the impact of common-shared regions on read distribution along transcript

The transcript-specific k-mer overlapping profile highlights an intrinsic relationship among related transcript isoforms that complicates the assignment of sequencing reads to individual isoforms. To visualize how common-shared regions affect read distribution along full-length transcripts, we analyzed both simulated and actual RNA-seq data by profiling consecutive 50-mer-associated counts. Our focus was on three inherently related isoforms: ACTB_ENST00000646664.1, ACTG1_ENST00000573283.7, and ACTG1P10_ENST00000428275.1 (as illustrated in Fig. 2a). Using a k-mer splitting strategy, we examined the global and homogeneous 50-mer overlapping profiles and compared these with 50-mer count profiles from both simulated and real sequencing data. Despite differences in overall sequence contexts, the global overlapping profile consistently showed a maximum overlap of 64 at a shared 50-mer position across all three isoforms. This 64-isoform-shared 50-mer region produced identical k-mer count peaks in the 50-mer count profiles of all three transcripts, whether derived from simulated or sequencing data. This finding suggests that the global overlapping pattern is closely aligned with the 50-mer counting pattern.

**Fig. 2.**
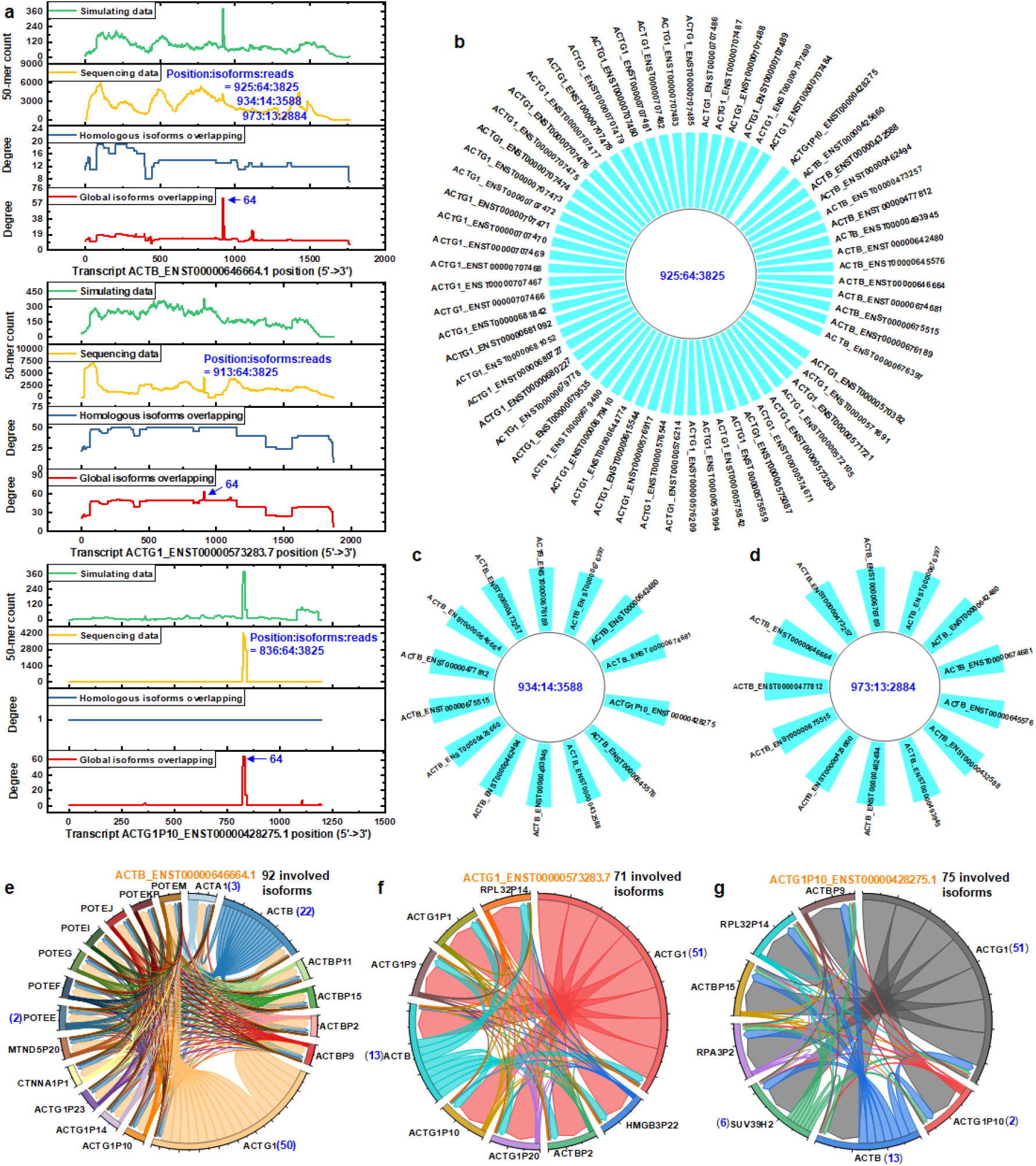
Analysis of common-shared regions among homologous and heterologous isoforms. **a**) A 50-mer sequence shared across 64 isoforms from three distinct genes. The analysis focuses on three isoforms-ACTB_ENST00000646664.1, ACTG1_ENST00000573283.7, and ACTG1P10_ENST00000428275.1-showing their homologous and global isoform overlap profiles based on the Homo_sapiens.GRCh38.cdna.all.fa reference. 50-mer counts were obtained from both polyester-simulated and empirical sequencing data. The common 50-mer sequence shared by all 64 isoforms is highlighted to indicate its presence in the three isoforms, with both sequencing and simulation data showing a prominent count peak in this shared region. **b**) A 50-mer located at position 925 within the ACTB_ENST00000646664.1 isoform is shared across 64 isoforms originating from three distinct genes. **c-d**) Two additional 50-mers, located at positions 934 and 973 within ACTB_ENST00000646664.1, are shared by 14 and 13 isoforms, respectively. **e**) The full set of 50-mers specific to ACTB_ENST00000646664.1 matches a total of 92 homologous and heterologous isoforms, spanning 19 different genes. **f**) The unique 50-mers associated with the isoform ACTG1_ENST00000573283.7 correspond to a set of 71 homologous and heterologous isoforms spanning 9 different genes. **g**) The 50-mers specific to ACTG1P10_ENST00000428275.1 are associated with 75 full set of homologous and heterologous isoforms, covering 8 different genes.

To provide a more detailed view of the global overlapping pattern within the individual isoform, we focused on three adjacent distinct overlapping points with degrees of 64, 14, and 13, corresponding to positions 925, 934, and 973 along ACTB_ENST00000646664.1 (as presented in Fig. 2a), respectively. At these positions, different sets of isoforms are involved, highlighting the complexity of accurately assigning reads to multiple overlapping isoforms. Each position is associated with a specific set of involved isoforms. For instance, at position 925 of ACTB_ENST00000646664.1, all 64 different isoforms from three heterologous genes share the same consecutive 50-mer set (as shown in Fig. 2b). This 50-mer is shared with 12 of the 23 homologous isoforms of the ACTB gene. At different positions, such as 934 and 973, the sets of isoforms sharing the common regions differ (as shown in Fig. 2c and 2d). This confirms that the specific k-mer sequence determines the specific set of involved isoforms. Since all three 50-mer positions are adjacent along the transcript reference, their sequences partially overlap. This adjacency results in partially overlapping sets of transcript isoforms, determined by the specific positions of each 50-mer sequence (as illustrated in Supporting Information Fig. S9). The overlap highlights that a one-base sliding k-mer splitting strategy effectively ensures each k-mer represents a distinct subsegment of the transcript sequence, while overlapping regions capture incremental sequence variations. This approach enables precise partitioning of the sequence into manageable units, maintaining continuity and preserving context for distinguishing isoform-specific or shared regions. In the context of complex 50-mer overlaps, these position-dependent k-mer-specific isoform sets significantly complicate read assignment for quantification. Matched reads must be accurately attributed to the correct isoforms, a task made more challenging by the involvement of both homologous and heterologous isoforms. The complexity arises from overlapping isoform sets at different positions, as matched reads corresponding to these 50-mers often intersect. Untangling these intricate relationships is exceedingly difficult, underscoring the need for highly precise methods to accurately distribute reads and resolve isoform-specific quantification.

By screening all k-mers along the transcript reference, we can enumerate all potential isoforms associated with the entire transcript, as each k-mer sequence maps to a specific set of isoforms. For instance, ACTB_ENST00000646664.1 is involved with 92 distinct isoforms derived from 19 heterologous genes (as illustrated in Fig. 2e). Among these, the ACTG1 gene contributes the largest number, with 50 isoforms. Although the transcript ACTG1_ENST00000573283.7 is slightly longer than ACTB_ENST00000646664.1, it is associated with fewer total isoforms-71 in total-and fewer involved genes-9 compared to ACTB’s 19 (as shown in Fig. 2f). This demonstrates that the diversity of involved isoforms is not necessarily correlated with the full length of the transcript. Furthermore, the shorter isoform ACTG1P10_ENST00000428275.1 is associated with 75 isoforms originating from 8 heterologous genes (as depicted in Fig. 2g), reinforcing the conclusion that transcript length does not directly influence isoform diversity.

### Validation of alignment ambiguity due to common-shared regions in RNA-seq

In RNA-seq analysis, alignment ambiguity often arises from common-shared regions between transcript isoforms. By splitting the transcript sequence into one-base sliding k-mers, we can theoretically construct a global overlapping profile specific to each transcript isoform. To validate the effect of these shared regions on ambiguous read assignments, we focus on a specific 59-nucleotide region common to 64 different isoforms across three heterologous genes. This region, highlighted in the ACTB_ENST00000646664.1 reference, contains 10 consecutive 50-mer sequences (as illustrated in Fig. 3a). The corresponding complementary DNA source is displayed at the genomic level based on the Homo_sapiens.GRCh38 reference. The highlighted 50-mer region corresponds to the regions (as shown in Fig. 2a). Notably, using long k-mer sequences can significantly increase the matching stringency between sequences, especially among heterologous genes. However, even with long k-mers, common shared regions are still widespread in transcript isoforms (as shown in Supporting Information Fig. S10). Specifically, the 500-mer sequence regions are shared among different heterologous genes, such as FAM89B (ENST00000449319.2) and ZNRT2 (ENST00000526433.1), both of which are protein-coding transcripts. To examine how shared regions influences the quantification of the involved isoforms using both alignment-based and alignment-free methods, we filtered the raw FastQ data by removing any reads containing this 59-nt sequence (10 consecutive 50-mer sequences). The 50-mer sequencing count profiles of ACTB_ENST00000646664.1, ACTG1_ENST00000573283.7, and ACTG1P10_ENST00000428275.1 were compared between the raw and filtered data (as shown in Fig. 3b). The removal of reads containing the 64-shared sequences led to a position-specific reduction in 50-mer counts across these isoforms. This pattern is expected across all 64 isoforms sharing the region.

**Fig. 3.**
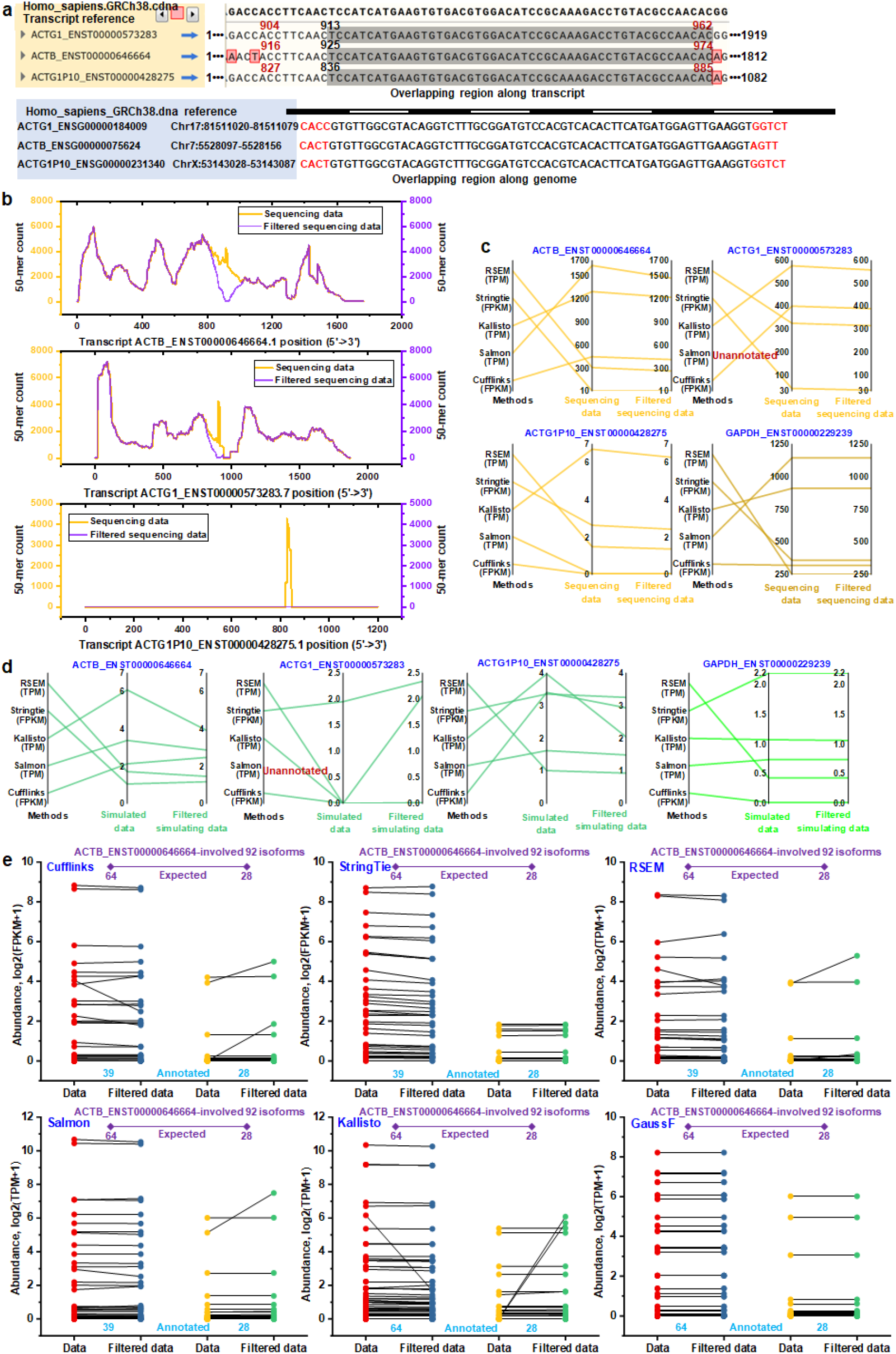
Validation of transcript isoform quantification impacted by shared regions. **a**) Overlapping regions at the transcript and gene reference levels: the shared region across three transcript isoforms-ACTB_ENST00000646664.1, ACTG1_ENST00000573283.7, and ACTG1P10_ENST00000428275.1-is aligned to their corresponding gene sequences. These regions are compared using the Homo_sapiens.GRCh38 reference genome. **b**) Sequencing count profiles of consecutive 50-mer along three representative isoforms, shown for both unfiltered (raw) and filtered datasets. Reads containing 50-mer sequences common to all 64 isoforms were excluded from both R1 and R2 sequencing datasets to create the filtered data. The sequencing count profile after filtering is contrasted with the original unfiltered data. **c**) The abundances of three isoforms involving a shared region and an unrelated isoform (GAPDH_ENST00000229239.10) were quantified using Cufflinks, Salmon, Kallisto, StringTie, and RSEM. Filtering out reads containing the shared region led to decreased abundance estimates for the involved isoforms across all quantification methods, whereas the abundance estimates for the unrelated isoform remained consistent between the original and filtered data. **d**) Abundance estimates for the three shared-region-involved isoforms and the unrelated isoform were obtained using Cufflinks, Salmon, Kallisto, StringTie, and RSEM on polyester-simulated datasets. The estimates for the shared-region-involved isoforms showed significant variation across different quantification methods, in contrast to the highly consistent estimates for the unrelated isoform. **e**) The quantification of all 92 isoforms involved within the entire ACTB_ENST00000646664.1 sequence is compared between the raw and filtered sequencing data. Of these, 64 isoforms share a sequence in the 916-974 region, while 28 share sequences in other regions. Using traditional methods (Cufflinks, Salmon, RSEM, and StringTie), 39 of the 64 isoforms were annotated, and the Kallisto method can identify all 64 isoforms. The abundance of the 64 isoforms generally decreases after filtering, while the 28 unrelated isoforms maintain consistent abundance across the methods. The transcript-specific minimal-shared region-based GaussF method shows consistent quantification for all 92 isoforms between the original and filtered data.

Beyond sequencing count profiles, we employed six quantification methods-Cufflinks, Salmon, Kallisto, StringTie, RSEM, and GaussF (a minimal-sharing strategy method)-to assess how the filtering affected isoform abundance (as seen in Fig. 3c). As a control, we quantified an unrelated isoform, GAPDH_ENST00000229239.10, which has no overlap with the 64-common-shared region. As expected, the abundance of the three involved isoforms showed a decreasing trend from the raw to filtered data across all quantification methods, reflecting the removal of related reads. In contrast, GAPDH abundance remained consistent between the raw and filtered data across all methods, validating that the common shared regions specifically correlate with the involved isoforms. We also explored the effects of common-shared regions using simulated data (as demonstrated in Fig. 3d). For this analysis, we set the initial isoform abundance ratio to 1 with a moderate total read count parameter in the Polyester package simulation. This setup intentionally kept the abundance of all isoforms relatively low, which was crucial for evaluating quantification accuracy across different methods. The implementation details for this simulation setup, including the code specifics for Polyester, are provided in the Methods section. By keeping the isoform abundance low, we aimed to amplify variability in quantification due to shared regions, allowing a clearer evaluation of their effects. We compared the 50-mer simulated count profiles of ACTB_ENST00000646664.1, ACTG1_ENST00000573283.7, and ACTG1P10_ENST00000428275.1 with their respective global, homologous, and local overlapping profiles between the raw and filtered data (as shown in Supporting Information Fig. S11). The removal of reads containing the 64-shared sequences resulted in position-specific reductions in 50-mer simulated counts across these isoforms, mirroring the observed sequencing count reduction trends. Interestingly, the simulated data did not consistently show the expected downward trend in isoform abundance after filtering. Instead, some quantification methods displayed an unexpected increase in abundance. However, the GAPDH control remained stable, reinforcing that the shared region was the primary cause of ambiguity. This inconsistency suggests that traditional quantification methods struggle with uncertain read assignments when confronted with shared regions, especially for the low-abundance transcript. While a decrease in abundance was anticipated in the filtered data, the observed increase in some cases suggests a more complex process of ambiguous read assignment, further complicating accurate quantification with traditional approaches.

To demonstrate how isoform overlapping patterns determine the distribution of sequencing reads along the transcript reference, we compared the global and homologous isoform 50-mer-based overlapping profiles of ACTB_ENST00000646664.1 and UBC_ENST00000339647.6 with their respective 50-mer sequencing counts across three different samples (as shown in Supporting Information Fig. S12). The consistent pattern of 50-mer sequencing counts observed at positions where the 64 isoforms share 50-mers within the ACTB_ENST00000646664.1 transcript reference aligns with the global overlapping pattern observed at these shared positions. In the UBC_ENST00000339647.6 comparison, a strong correlation between position-specific 50-mer sequencing counts and the global overlapping profile was observed (as shown in Supporting Information Fig. S12, lower panel), highlighting how complex overlaps between heterologous genes reshape sequencing count profiles. Additionally, the 50-mer sequencing count profiles for many other individual transcripts-HPRT1_ENST00000298556.8, B2M_ENST00000349264.10, RPLP0_ENST00000392514.9, and TUBB_ENST00000383564.8-remained highly consistent across different samples (as illustrated in Supporting Information Fig. S13). The reproducibility of sequencing profiles across transcripts and samples supports the notion that k-mer and read distribution patterns are likely predefined and predictable, rather than random or variable.

To further investigate, we expanded our analysis of sequencing raw and filtered data to all 92 isoforms related to ACTB_ENST00000646664.1, including the 64 isoforms associated with the 916-974 common-shared region and 28 isoforms that do not share this region but are instead linked to other regions (as illustrated in Fig. 3e). Using the same six quantification methods, we found that Cufflinks, Salmon, StringTie, and RSEM annotated only 39 of the expected 64 isoforms, while Kallisto and GaussF successfully identified all 64. Notably, “annotated” transcripts refer to those present in the reference list for quantification, while “detected” transcripts correspond to those with measurable abundance from the selected reference transcripts. In the set of 64 isoforms (or the subset of 39 annotated by certain methods), the general trend showed a decrease in isoform abundance from raw to filtered data. However, many exceptions were observed with traditional methods, where certain isoforms demonstrated either consistent or elevated abundance. In contrast, the GaussF method, which focuses on minimal-shared regions, demonstrated a high level of consistency, maintaining same abundance levels between raw and filtered data. Since the 64-isoform common-shared region does not overlap with the minimal-shared region utilized by GaussF, its abundance quantification is unaffected. Additionally, the set of isoforms identified across the five quantification methods-Cufflinks, Salmon, StringTie, RSEM, and GaussF (comparable to Kallisto)-were analyzed to identify method-specific isoform sets (as shown in Supporting Information Fig. S14). Although Cufflinks, Salmon, StringTie, and RSEM each annotated 39 isoforms, the isoform sets from Cufflinks, StringTie, and RSEM were identical. In contrast, only 14 of the 39 isoforms annotated by Salmon overlapped with those identified by Cufflinks, StringTie, and RSEM. This variation in method-determined isoform sets highlights the potential discrepancies in transcript profiles produced by different quantification methods, which can, in turn, lead to differing interpretations in downstream biological function analyses. For the 28 isoforms unrelated to the 916-974 region, the majority displayed consistent abundance between the raw and filtered data, which aligns with expectations. However, discrepancies emerged when comparing quantification across different methods such as Cufflinks, Salmon, Kallisto, and RSEM. These methods showed an increased abundance for several isoforms, suggesting complex ambiguity in read assignment. This observation underscores the importance of considering read ambiguity in RNA-seq quantification, particularly when analyzing complex transcript structures with overlapping regions. The inability to trace the exact origin of ambiguous reads in probabilistic (likelihood) models highlights the inherent challenges in accurately assigning reads to specific isoforms. This limitation emphasizes the need for careful interpretation of quantification results. The GaussF method stands out in this context by maintaining consistent isoform abundance through its minimal-shared region approach. This strategy effectively mitigates the confounding effects of shared sequences, offering a robust solution for RNA-seq quantification. The minimal-shared region ensures that read assignments are less influenced by overlapping sequences, resulting in more reliable quantification even in complex genomic contexts.

### Overlapping analysis of sequences across varying k-mer lengths in multiple species

Starting with human transcripts, we employ a k-mer splitting strategy, which involves dividing each transcript sequence into a set of k-mers. Each k-mer in this set is then matched against k-mers from other transcripts to construct a comprehensive k-mer overlapping profile (as illustrated in Fig. 1d and 1h). The choice of k-mer length significantly influences the resolution of these profiles and the stringency of the matching process. By varying k-mer lengths-such as 50, 100, 250, and 500-the characteristics of the overlapping profiles can be altered. This effect is exemplified in the ACTB_ENST00000646664.1 transcript sequence (as demonstrated in Fig. 4a). Adjusting k-mer length provides a trade-off between specificity and sensitivity in genomic analyses. Longer k-mers offer higher specificity by demanding more stringent sequence matches, thereby reducing the cumulative count of heterologous isoforms and the maximum overlap degree. This is because the stricter matching conditions associated with longer k-mers limit the identification of shorter, less specific overlaps. Conversely, shorter k-mers enhance sensitivity to shared regions, capturing a broader range of overlaps but often at the cost of increased computational complexity and potentially higher false-positive rates. At a broader genomic scale, the analysis of common-shared regions using k-mer lengths of 50, 100, 250, and 500 reveals that longer k-mers lead to a significant reduction in shared regions across genes. Here, each gene sequence is defined by the combination of individual homologous transcript isoform sequences. For instance, at a 50-mer length, 62.4% of individual genes share at least one 50-mer sequence with other genes. However, this percentage decreases to 25.8% with 500-mer sequences. In this case, the highest overlap degree (45) is observed in genes such as DUX4, while the average overlap degree is around 7.1 (see Fig. 4b and 4c). This trend implies that the larger the k-mer, the less frequent the overlaps, reducing the complexity of distinguishing specific gene features but also diminishing the granularity of the analysis due to substantial overlapping regions.

**Fig. 4.**
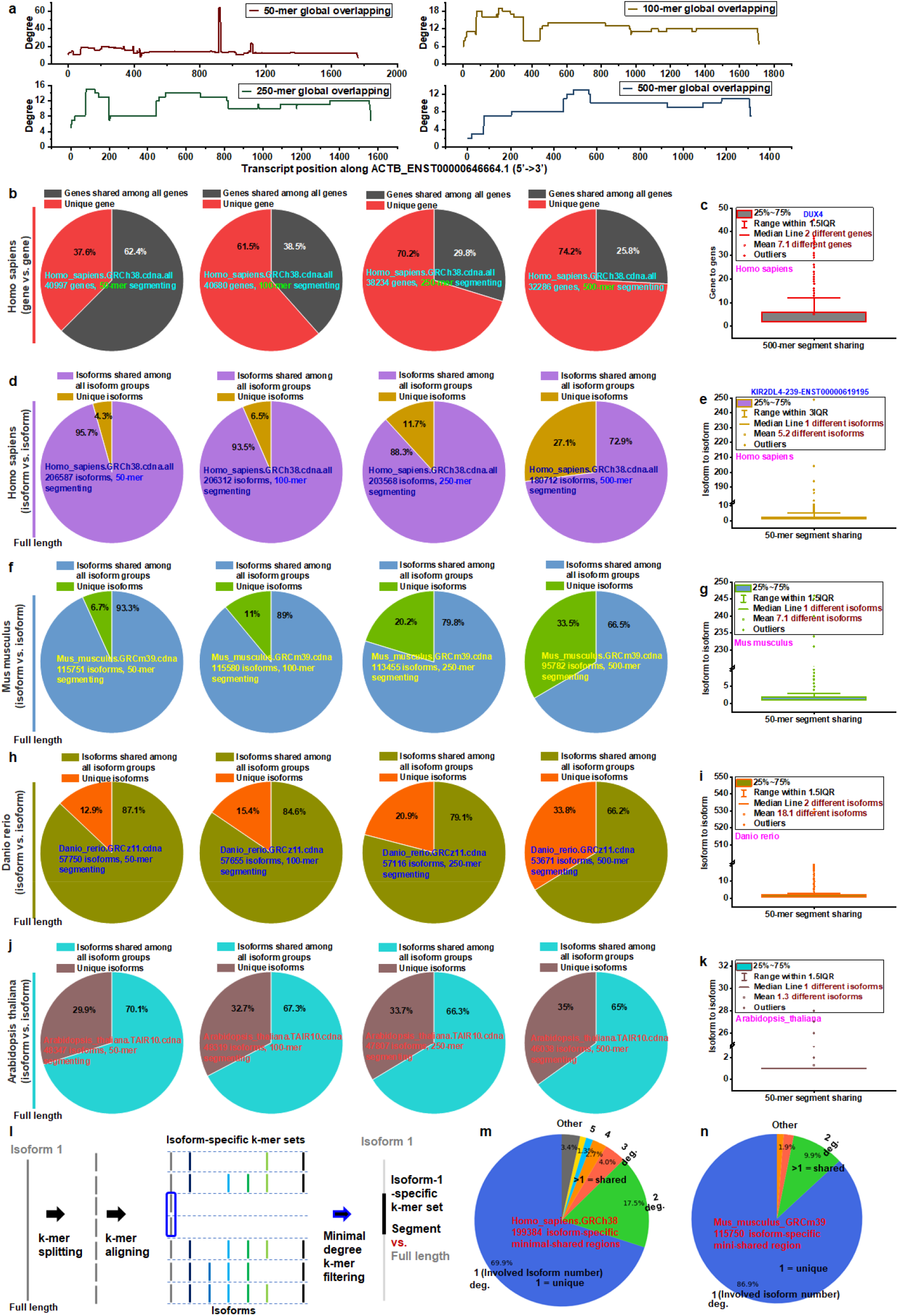
Overlapping analysis of sequences across varying k-mer lengths in multiple species. **a**) Global overlapping profile of Homo sapiens ACTB_ENST00000646664.1, generated by analyzing common-shared regions using different k-mer lengths (50-mer, 100-mer, 250-mer, and 500-mer). **b**) Overlapping analysis at the gene level in Homo sapiens across k-mer lengths from 50 to 500 nucleotides. At the 50-mer level, 37.5% of genes contain a unique set of sequences. This proportion increases to 72.2% at the 500-mer level, indicating that as k-mer length increases, more genes are characterized by unique sequences. **c**) At the 500-mer level, each gene shares its entire set of 500-mers with an average of 7 to 8 other genes. Notably, at the 50-mer level, the DUX4 gene (ENSG00000283949) shares its entire set of 500-mers with 44 other genes. **d**) Overlap analysis at the Homo sapiens transcript isoform level indicates that only 4.3% of isoforms contain a unique 50-mer sequence set. This proportion increases significantly to 27.1% at the 500-mer level, suggesting greater specificity at longer sequence lengths. **e**) At the 50-mer level, each Homo sapiens isoform shares its entire 50-mer sequence set with an average of 5–6 other isoforms. The isoform ENST00000672643.1 (gene KIR2DL4) shares its 50-mer segment with a total of 249 other transcript isoforms. **f**) In Mus musculus (GRCm39), 6.7% of isoforms have a unique 50-mer sequence, which increases to 33.5% at the 500-mer level. **g**) At the 50-mer level, each Mus musculus isoform shares its sequence set with an average of 7-8 other isoforms, with a maximum of 246 common-shared isoforms. **h**) In Danio rerio (GRCz11), 12.9% of isoforms are specified by a unique 50-mer sequence, increasing to 33.8% at the 500-mer level. **i**) On average, each Danio rerio isoform at the 50-mer level shares its sequence set with 18-19 other isoforms, with the maximum shared instance involving 531 isoforms. **j**) In Arabidopsis thaliana (TAIR10), 29.9% of isoforms have a unique 50-mer sequence, rising to 35% at the 500-mer level. **k**) At the 50-mer level, each Arabidopsis thaliana isoform shares its sequence with an average of 1-2 other isoforms, with a maximum shared sequence involving 28 isoforms. **l**) Minimal-shared regions identified through k-mer alignment enable the isolation of isoform-specific regions, enhancing the precision of isoform characterization. These minimal-shared segments are shorter than the entire reference isoform sequence. **m**) For Homo sapiens, the isoform-specific minimal-shared regions gene symbol-annotated 199,384 isoforms. Of these, 69.9% have a unique minimal-shared region (degree 1), meaning they are exclusive to a single isoform. A degree higher than 1 indicates that the minimal-shared region is shared by multiple isoforms, meaning no unique region exists specifically for the isoform. Well-annotated isoforms undergo k-mer splitting during minimal-shared filtering to enhance specificity. **n**) For Mus musculus, the isoform-specific minimal-shared regions annotated 115,750 isoforms. Here, 86.9% of isoforms have a unique minimal-shared region (degree 1). A degree higher than 1 signifies that the region is shared with other isoforms, indicating no unique region exists specifically for that isoform. To increase specificity, well-annotated isoforms undergo k-mer splitting during minimal-shared filtering.

Expanding k-mer analysis to the human transcriptomic level allows for the segmentation of each transcript into distinct k-mer sets, enabling a more refined resolution of overlapping profiles and revealing isoform-specific patterns unique to each transcript. The length of the k-mer remains a critical factor, influencing the specificity and granularity of these isoform patterns. This approach provides a more nuanced understanding of transcript diversity and highlights subtle variations between isoforms. For instance, with a k-mer length of 50, only 4.3% of transcript isoforms maintain high sequence uniqueness without overlapping other isoforms. However, as k-mer length rises to 500, the proportion of unique isoforms increases to 27.1% (as depicted in Fig. 4d). This increase is likely due to the longer k-mer segments imposing more stringent matching conditions. Additionally, the total isoform count decreases as k-mer length increases, since transcripts shorter than 500 nt are excluded from the 500-mer analysis. Aggregating all isoforms from the shared isoforms set, the mean count of isoforms that share sequences with a given isoform is approximately 5-6 (as illustrated in Fig. 4e). However, some isoforms show exceptionally high overlap, such as ENST00000672643.1 (KIR2DL4), which shares a 50-mer segment with 249 other transcript isoforms from heterologous genes (as shown in Supporting Information Fig. S15). The KIR (Killer-cell Immunoglobulin-like Receptor) genes play a crucial role in the immune system.[30] They encode proteins that interact with MHC class I molecules (Major Histocompatibility Complex class I) on other cells, enabling natural killer (NK) cells to distinguish between healthy and abnormal cells. Given the significant variability in KIR gene configurations across individuals,[31] the observed high degree of overlap is consistent with this genetic diversity. Further, across k-mer lengths from 50 to 500 nucleotides, we examined the regions shared between transcript isoforms and other heterologous genes (isoforms) to understand the contribution of involved genes across all isoforms (as shown in Supporting Information Fig. S16). Comparing overlapping isoforms based on k-mer length, we find that the proportion of unique isoforms, which do not share any 50-mer region with other heterologous isoforms, rises from 4.3% (as depicted in Fig. 4d) to 69.9%. For the 500-mer regions, this increase is even more substantial, from 27.1% to 94%. This suggests that most shared regions are primarily derived from homologous isoforms, with heterologous overlaps decreasing significantly as k-mer length increases.

To further clarify the distribution of common-shared isoforms across species, we applied k-mer splitting to the isoforms of Mus musculus for comparative analysis. The overall overlapping pattern in Mus musculus shows a similar trend to that of Homo sapiens, with the percentage of shared isoforms among all isoforms changing in a comparable manner (as depicted in Fig. 4f). Specifically, the percentage of unique isoforms increases as the k-mer length extends from 50 to 500. On average, each common-shared isoform in Mus musculus shares 50-mer sequences with 7 to 8 other isoforms (as illustrated in Fig. 4g). For Danio rerio, the percentage of unique isoforms rises from 12.9% to 33.8% as the k-mer length increases from 50 to 500 (as demonstrated in Fig. 4h). Each common-shared isoform in this species shares 50-mer regions with approximately 18 to 19 other isoforms on average (as presented in Fig. 4i). In Homo sapiens, Mus musculus, and Danio rerio, there is a significant increase in the percentage of unique isoforms as the k-mer length grows from 50 to 500. In contrast, the plant Arabidopsis thaliana exhibits a different pattern. The percentage of unique isoforms only slightly increases from 29.9% to 35% as the k-mer length extends (as highlighted in Fig. 4j). Moreover, each common-shared isoform in Arabidopsis shares 50-mer regions with only 1 to 2 other isoforms (as depicted in Fig. 4k). This distinct pattern in the plant species differs notably from those observed in the aforementioned animal species.

To improve isoform quantification in RNA-seq analysis across species, we employ a minimal-sharing strategy to filter out regions with extensive overlap, retaining only minimally shared regions to represent each isoform (as shown in Fig. 4l). This approach focuses on unique or nearly unique regions of each isoform, even within complex and overlapping transcript sequences. For each isoform, the minimal-shared degree across k-mers may vary: a degree of 1 indicates the presence of unique regions specific to that isoform, while a higher degree signifies regions shared by multiple isoforms. For instance, a degree of 2 indicates that a region is shared by at least two isoforms. Because existing algorithms struggle to allocate reads from highly overlapping regions to specific isoforms with high confidence, we define quantification based on minimal-shared regions as an aggregate contribution from all involved isoforms, thereby avoiding uncertain assignments. In Homo sapiens, 69.9% of well-annotated isoforms contain unique regions based on the 50-mer segmentation approach (as depicted in Fig. 4m), while 17.5% exhibit a minimal-shared degree of 2, indicating that two isoforms contribute to the quantification. In Mus musculus, 86.9% of isoforms are characterized by minimally shared, unique regions (degree 1), and 9.9% have a minimal-shared degree of 2 (as shown in Fig. 4n). For isoforms with a minimal-shared degree greater than 1, downstream analyses should clearly indicate the aggregate nature of quantification to ensure accurate data interpretation and prevent erroneous assignments.

### Validating minimal-sharing strategies for enhanced transcriptome quantification in a mouse model of Huntington’s disease

The minimal-sharing strategy leverages distinct, minimally shared regions within entire transcript sequences to characterize specific isoforms. This approach enables us to understand how these targeted regions, rather than full-length transcripts, can represent biological processes across the transcriptome. To examine the effects of Huntington’s disease (HD) on transcript expression, we conducted RNA-seq analysis on three HD mouse models and their corresponding wild-type (WT) controls. We compared the transcripts detected in HD and WT samples using six quantification methods: Cufflinks (148,980 reference transcripts), StringTie (149,139 reference transcripts), Salmon (115,300 reference transcripts), RSEM (149,139 reference transcripts), Kallisto (115,989 reference transcripts), and GaussF (115,751 reference transcripts), as shown in Figure 5a. (as depicted in Fig. 5a). Each method utilized its own specific reference transcript set for mapping and quantification. Ideally, the set of detected isoforms that differ between HD and WT samples should highlight distinct transcriptomic features directly associated with Huntington’s disease. However, our results reveal that the degree of overlap varies significantly by quantification method, with Cufflinks identifying the highest number of isoforms and overlapped isoforms compared to other approaches. This suggests that the choice of quantification method can substantially impact the set of detectable isoforms and, consequently, interpretations of disease-associated transcriptomic differences. To further examine the internal variation in abundance estimations, we performed principal component analysis (PCA) using the transcriptomic abundance profiles obtained from the different methods (as shown in Fig. 5b). The PCA based on GaussF quantification revealed that the first principal component (PC1) explained 89% of the variance, while the second principal component (PC2) accounted for only 4%. This result contrasts with those from other methods: Salmon (PC1: 68%, PC2: 9%), RSEM (PC1: 63%, PC2: 10%), Kallisto (PC1: 68%, PC2: 11%), Cufflinks (PC1: 39%, PC2: 25%), and StringTie (PC1: 26%, PC2: 22%). The genomic alignment-based methods, Cufflinks and StringTie, showed a more even distribution of variance across the first two principal components, suggesting a more complex or multidimensional data structure. In contrast, the alignment-free methods indicated a single dominant source of variation driving the data structure. Specifically, the GaussF-based PCA suggests that most of the variance can be captured by PC1 alone, indicating a strong underlying trend or pattern in the transcript abundance dataset. Considering that the HD samples are characterized by a single htt gene mutation compared to the WT samples, the minimal shared region-based GaussF method appears to capture this primary source of variation effectively. This association underscores the effectiveness of using minimal shared regions in capturing significant biological variation at the transcriptome level.

**Fig. 5.**
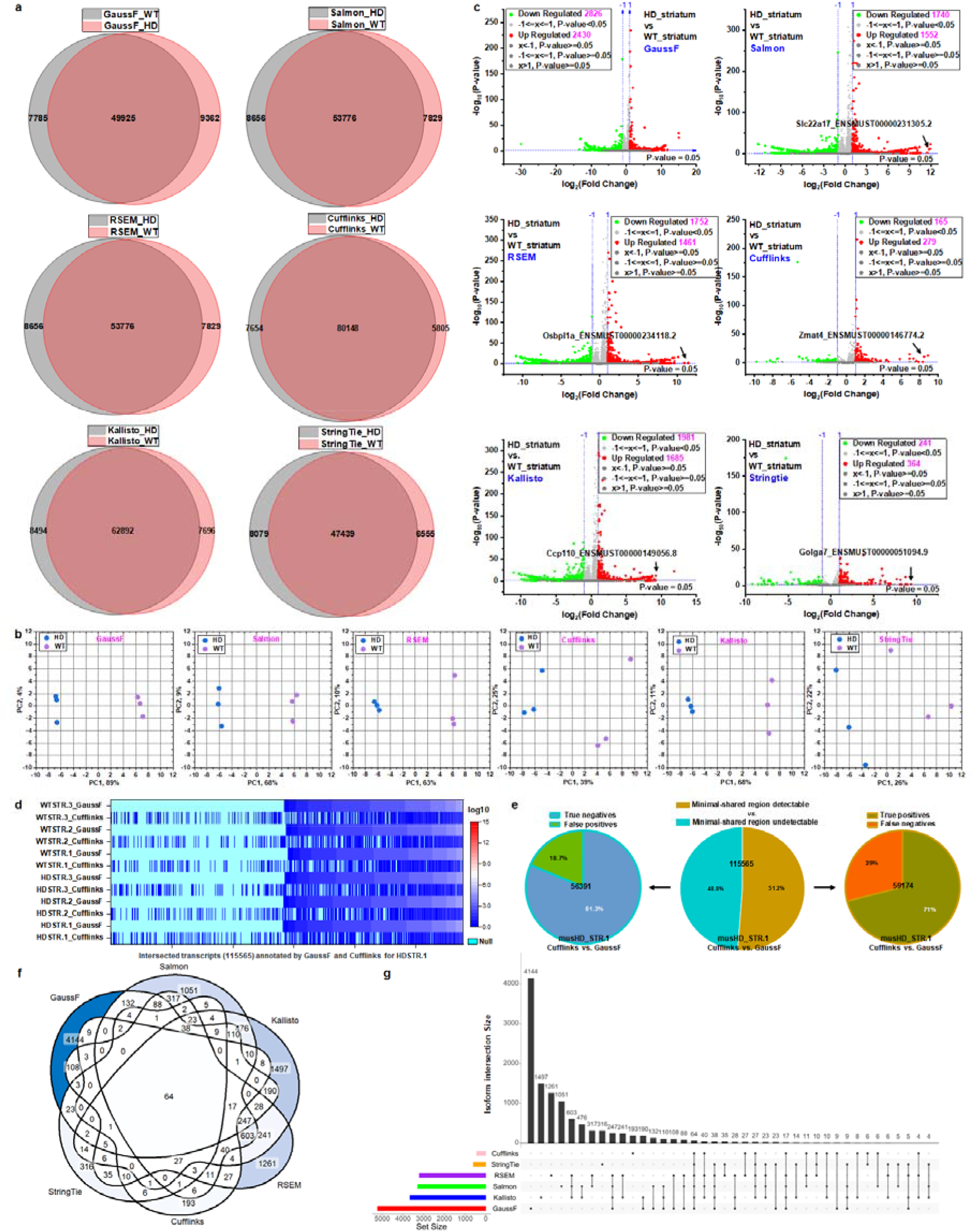
Validation of minimal-sharing strategy-based transcriptome quantification. **a**) Venn diagram illustrating the differences in the number of detectable isoforms from RNA-seq data of mouse Huntington’s Disease model (HD) and wild-type (WT) samples, quantified using six methods: Cufflinks, StringTie, Salmon, RSEM, Kallisto, and GaussF (featured by minimal-shared region). **b**) Principal Component Analysis (PCA) of gene expression at transcript isoform level data from six samples, comparing musHD (n = 3) and musWT (n = 3). The PCA plot displays the first two principal components (PC1 and PC2), which account for the largest variance in the data. The X-axis (PC1) explains the majority of the variance, while the Y-axis (PC2) explains the next largest variance. **c**) Differential gene expression analysis between HD (n = 3) and WT (n = 3), based on transcript count profiles obtained from different quantification methods. The plot shows log2 fold changes on the X-axis and the -log10 p-values on the Y-axis. Each point represents a transcript isoform, with significant differentially expressed isoforms (p-value < 0.05 and |log2 fold change| > 1) highlighted. **d**) Heatmap of transcript abundances for annotated transcripts common between GaussF and Cufflinks quantifications, across three HD and three WT samples. Transcripts are sorted based on their abundance rankings determined by GaussF in the HD STR-1 sample, allowing for direct comparison with the corresponding abundance values determined by Cufflinks. **e**) Comparison of annotated transcript detection for the musHD_STR-1 sample. All union set of transcripts (115565) were grouped into two sets based on minimal shared regions: detectable transcripts (59174) and undetectable transcripts (56391) determined by GaussF. Among the undetectable group, 18.7% are identified by Cufflinks as false positives. In contrast, within the detectable group, 29% are missed by Cufflinks and classified as false negatives. **f**) Venn diagram showing the overlap of significant differentially expressed isoforms across the six quantification methods. **g)** Upset plot displaying the intersections in detail of significant differentially expressed isoforms identified by the six methods. Vertical bars represent the number of isoforms shared by specific combinations of methods, as indicated by the connected dots below. The horizontal bar chart on the left shows the total number of significant isoforms identified by each individual method. All possible combinations of overlapping isoforms between methods are presented.

Building on these findings, we conducted Differential Gene Expression Analysis (DGEA) of the sequencing count profiles from HD and WT samples-a key step in downstream biological analysis. Analyzing the same raw sequencing datasets (as shown in Fig. 5c), different quantification methods produced varying differential gene expression patterns, indicating that expression results depend on the quantification method used. For instance, although Cufflinks identified the highest number of detectable isoforms (see Fig. 5a), it reported the lowest total number of significantly downregulated (165) and upregulated (279) isoforms in the DGEA compared to other methods. Similarly, the genomic alignment-based method StringTie produced a low number of differentially expressed isoforms comparable to Cufflinks. This highlights the impact of quantification tools on DGEA outcomes. The probabilistic models underlying genomic alignment-based methods may accommodate a broader range of transcripts due to widespread soft clipping events, potentially reducing sensitivity in detecting abundance changes. In contrast, genomic alignment-free methods like Salmon, Kallisto, and RSEM identified a higher number of significant isoforms. Notably, the GaussF method-which relies on minimal shared regions and identified the highest number of differentially expressed isoforms-enhanced the contrast between HD and WT samples by eliminating ambiguity during quantification. This minimal-sharing strategy likely improves both the sensitivity and stringency in declaring isoforms as differentially expressed, effectively capturing significant biological variations between the HD and WT samples.

To validate the effectiveness of the minimal-shared region approach in GaussF compared to traditional methods, we examined upregulated transcripts in WT identified differently by individual method (see Fig. 5c). These transcripts, highlighted by different methods, exhibit some of the highest log-fold changes with statistically significant differences. For instance, the RSEM method identifies Osbpl1a_ENSMUST00000234118.2 as significantly downregulated in HD samples compared to WT samples. To investigate the read-mapping coverage of this transcript, we profiled the 50-mer sequencing counts and global overlapping degree across the full transcript sequence in three HD and three WT samples (as shown Supporting Information Fig. S17). The overlapping profile indicates extensive isoform overlap with homologous and heterologous genes, contributing to mapping ambiguity. While RSEM detects high read-count abundance for this transcript in two of the three WT samples, showing significant differences among them, a closer examination of the zoomed-in 50-mer coverage reveals no substantial differences between the WT samples. This discrepancy is likely due to probabilistic read assignment. Furthermore, the Kallisto method presented clear instances of false positives (as depicted in Supporting Information Fig. S18). For example, Ccp110_ENSMUST00000149056.8 was detected by Kallisto with high read abundance in three WT samples; however, zoomed-in 50-mer coverage reveals that no mapped 50-mer sequences from the entire sequencing datasets actually align to the minimal-shared regions in WT samples, indicating that detection of this transcript is improbable. In addition to false positives, false negatives were observed with the Salmon method (as shown in Supporting Information Fig. S19). For Slc22a17_ENSMUST00000231305.2, Salmon detected null abundance in all three HD samples, identifying it as downregulated. Yet, 50-mer sequencing coverage clearly shows reads aligning to the minimal-shared regions, contributing to transcript abundance. Salmon’s probabilistic model appears unable to capture this alignment, resulting in a false negative. Similar false negatives were observed with Cufflinks and StringTie for Zmat4_ENSMUST00000146774.2 and Golga7_ENSMUST00000051094.9 (as demonstrated Supporting Information Fig. S20 and Fig. S21). Moreover, traditional methods reported approximately 1,000-fold higher abundance differences for five specific transcripts between WT and HD samples. However, 50-mer sequencing count profiles across three WT and three HD samples did not consistently support such large differences, revealing inaccuracies in these methods. To evaluate transcriptome-wide performance, we used GaussF-which leverages minimal-shared regions-as the ground truth for identifying false positives and false negatives. By definition, if sequencing read segments align to a transcript’s minimal-shared regions, the presence of transcript-specific templated reads in the sequencing data ensures that the transcript should be detectable by any method. Conversely, if no sequencing read segments align to these transcript-specific minimal-shared regions, the absence of corresponding reads indicates that the transcript should not be detected. Errors in read assignment can result in false positives-transcripts incorrectly identified as present-or false negatives-transcripts that are missed despite being detectable. A comparison between GaussF and Cufflinks revealed a significant number of both false positives and false negatives (as shown in Fig. 5d), emphasizing the limitations of traditional quantification approaches. In the HD_STR.1 sample, we categorized the union set of transcripts detected by GaussF and Cufflinks into two groups based on GaussF’s minimal-shared regions: detectable and undetectable. Among the undetectable transcripts (as shown in Fig. 5e), 18.7% were erroneously reported as present by Cufflinks (false positives). Despite the complete absence of sequencing reads or read segments aligning to these transcripts’ minimal-shared regions, Cufflinks assigned them high abundance counts. For instance, the 50-mer profiles of transcripts such as ENSMUST00000160682, ENSMUST00000231569, ENSMUST00000080813, ENSMUST00000135134, ENSMUST00000119726, and so on, showed no counts aligning to their transcript-specific minimal-shared regions, yet Cufflinks detected them with inflated abundance (as detailed in Supporting Information Fig. S22). On the other hand, among the detectable transcripts, 29% were missed by Cufflinks (false negatives) (as shown in Fig. 5e). In these cases, sequencing read segments clearly aligned to the transcript-specific minimal-shared regions-often at a unique alignment (as illustrated in Supporting Information Fig. S23)-yet Cufflinks failed to detect these transcripts, reporting null outputs despite the presence of unambiguous evidence. This discrepancy underscores how Cufflinks struggles to accurately identify transcripts even when sequencing data aligns to their unique regions. Notably, most minimal shared regions correspond to a minimal overlapping degree of 1. However, some regions exhibit a minimal shared degree greater than 1 (Fig. 4n), where sequencing counts could be distributed across all involved transcripts. Transcripts lacking unique regions (minimal shared degree = 1) are difficult to resolve at the isoform level. Thus, we classify such transcripts with a minimal shared degree exceeding 1 as false negatives in this context. These findings underscore the inherent limitations of traditional quantification methods, which rely on probabilistic models and are prone to ambiguities in read assignment. In contrast, GaussF quantifies transcript abundance with high specificity and sensitivity by focusing on minimal-shared regions. This approach eliminates false positives and false negatives, providing a reliable and biologically accurate transcript profile.

A further comparison of the differentially expressed isoform sets across six methods was conducted to analyze the impact of each method on identifying differentially expressed isoforms. The overlap of isoform sets from the six methods (as illustrated in Fig. 5f) reveals that only 64 isoforms are commonly shared, a number much lower than the total differentially expressed isoforms identified by any individual method. This observation suggests that the differential expression patterns are heavily influenced by the quantification method used. A more detailed dissection of the overlapping patterns (as shown in Fig. 5g) indicates the presence of method-specific isoform sets, especially in genomic alignment-free methods such as GaussF, Kallisto, Salmon, and RSEM. In contrast, genomic alignment-based methods like Cufflinks and StringTie also exhibit unique isoform sets, but the majority of isoforms identified by these methods are shared with those determined by other methods. These results confirm that differences in quantification models-particularly in how reads are probabilistically assigned-can lead to distinct transcriptome profiles and unique downstream patterns of differentially expressed isoforms.

### Enhanced functional insights into Huntington’s disease via comparative transcript quantification in a mouse model

To gain deeper functional insights into HD, we examined the unique characteristics of the HD model by highlighting key biological functions revealed through different quantification profiles. Utilizing Gene Ontology (GO) enrichment analysis-which provides a detailed classification of gene functions-we identified significant biological processes from an extensive set of GO terms, particularly those highly specific to isoform expression differences in transcriptome profiles. In our comparison of GO analysis results for biological processes, we utilized six quantification methods, including GaussF and Cufflinks (as illustrated in Fig. 6a) and StringTie, RSEM, Salmon, and Kallisto (as shown in supporting information Fig. S24). We identified differentially expressed isoforms using a p-value threshold of less than 0.05, without imposing fold-change requirements. Consistently across all six quantification methods, the primary biological process identified was related to the generation of precursor metabolites and energy-aligning with known features of HD.[32] However, the subsequent top nine biological processes varied among the methods. The GaussF method revealed a broader diversity of enriched biological processes within the top ten. In contrast, other methods prominently featured mitochondrial dysfunction and energy metabolism abnormalities among their top ten processes, whereas these appeared outside the top ten with GaussF. To further assess GaussF’s ability to capture key features of HD, we performed Kyoto Encyclopedia of Genes and Genomes (KEGG) pathway analysis, which offers a higher-level and comprehensive overview by examining interactions among multiple genes. Unlike the more specific GO terms, KEGG pathways are broader and aggregate functions into larger categories. The transcriptome profiling using GaussF highlighted significant enrichment in several neurodegenerative diseases-including HD, Parkinson’s Disease (PD), and Alzheimer’s Disease (AD) (as shown in Figure 6b).[33] Similar KEGG pathway enrichment patterns were observed with other quantification methods (detailed in Supporting Information Figure S25), reinforcing the consistency of these findings. This consistency underscores GaussF’s effectiveness in capturing the fundamental features of HD.

**Fig. 6.**
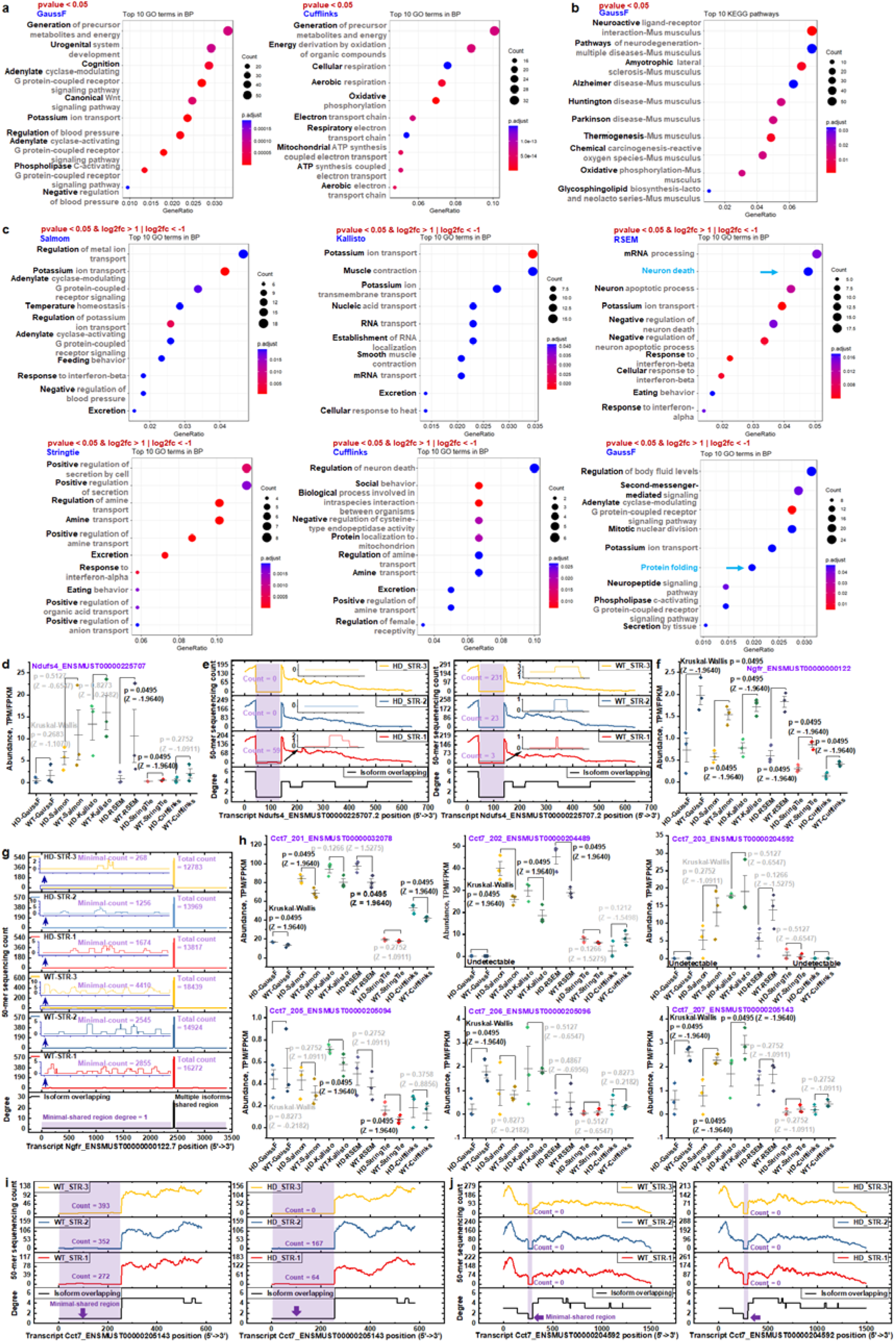
A detailed analysis of transcript isoforms assigned to specific biological functions. **a**) Comparative analysis of the top 10 enriched GO biological processes identified using GaussF’s minimal-sharing strategy and the traditional Cufflinks approach. Enrichment significance was determined for transcript isoforms with p-values <0.05, with processes ranked by significance and count. **b**) Top 10 KEGG pathways identified based on transcriptome profiles generated by GaussF, corresponding to the GO biological processes in panel (a). Pathways are ranked by enrichment significance and count for input isoforms with p-values <0.05. **c**) GO biological process enrichment analysis for transcript isoforms identified with p-values <0.05 and log2 fold changes >1 or <−1 across six quantification methods: Cufflinks, StringTie, Salmon, RSEM, Kallisto, and GaussF (minimal-sharing strategy). This comparison highlights method-dependent variations in biological process enrichment. **d**) RSEM-based GO biological process analysis identifies neuron death involvement, highlighting the Ndufs4 gene. Isoform abundance for Ndufs4_ENSMUST00000225707.2 is compared between HD (n=3) and WT (n=3) samples across the six quantification methods. **e**) 50-mer isoform overlap degree and sequencing count profile for Ndufs4_ENSMUST00000225707.2, with a zoomed-in view of the minimal-shared region and its corresponding sequencing count profile. **f**) Abundance estimates for neuron death-related gene Ngfr_ENSMUST00000000122.7 in HD and WT samples across six quantification methods. **g**) 50-mer isoform overlap degree and sequencing count profile for Ngfr_ENSMUST00000000122.7, with an inset focusing on the minimal-shared region and sequencing count profile. The total count and minimal count are represented as the 50-mer count sums across the entire transcript and the minimal-shared region, respectively. **h**) GaussF-based GO biological process analysis identifies the protein folding process involving the Cct7 gene. Abundance estimates are shown for six detected isoforms-Cct7_ENSMUST00000032078.9, Cct7_ENSMUST00000204489.3, Cct7_ENSMUST00000204592.3, Cct7_ENSMUST00000205094.2, Cct7_ENSMUST00000205096.2, and Cct7_ENSMUST00000205143.2-across the six quantification methods. **i**) 50-mer overlap and sequencing count profile for Cct7_ENSMUST00000205143.2, with a focus on the minimal-shared region. **j**) 50-mer overlap and sequencing count profile for Cct7_ENSMUST00000204592.3, with a focus on the minimal-shared region. All RNA-seq data were generated from HD (n=3) and WT (n=3) samples. Population differences were evaluated using a non-parametric Kruskal-Wallis test.

When considering all transcripts with a statistically significant p-value (e.g., p < 0.05) without applying a log fold change (logFC) threshold, these transcripts pass the significance test, the magnitude of their expression changes may be minimal and possibly not biologically meaningful. Therefore, to focus on transcripts with both statistically significant and biologically meaningful changes, we set a p-value threshold of < 0.05 and applied a logFC threshold of > 1 or < –1 to filter isoforms for GO analysis. Under these conditions, we conducted GO analyses based on transcript profiles derived from six quantification methods (as shown in Fig. 6c). Incorporating a logFC threshold alongside the p-value cutoff further refines the analysis by filtering out transcripts with minimal expression changes, thereby enhancing the specificity and depth of the insights gained. This dual-threshold approach enables a more focused identification of biologically significant transcripts, allowing for a clearer understanding of differential expression patterns in the HD model and improving the interpretability of the GO enrichment results. This adjustment reshapes the patterns of GO biological processes across the different quantification methods. There is no consistent primary biological process identified across all six methods. The number of genes corresponding to the top 10 biological process items is significantly lower than when including all statistically significant transcripts without a logFC threshold (as shown in Fig. 6a), especially in the GO analyses based on StringTie and Cufflinks. The maximum gene counts for the primary biological processes are 8 and 6 for StringTie and Cufflinks, respectively, aligning with the low number of differentially expressed isoforms these methods identified (as shown in Fig. 5c). These filtered transcript sets theoretically provide more biologically insightful and interpretable results for HD. In the RSEM-based biological process analysis, neurodegeneration-related neuronal death processes are enriched, as highlighted. HD is a neurodegenerative disorder characterized by the progressive loss of neurons, particularly in the striatum and cortex regions of the brain.[34] In the GaussF-based biological process analysis, the enrichment of the protein folding process is noteworthy, as protein misfolding is a primary event in HD pathogenesis.[35]

To explore how differentially expressed genes contribute to specific biological processes, we investigated the assignment of differential expression to processes such as neuron death using RSEM (as shown in Fig. 6c). Focusing on the neuron death-associated gene Ndufs4, we analyzed all six of its transcript isoforms, including Ndufs4_ENSMUST00000225707.2 (as shown in Fig. 6d) and others (as listed in Supporting Information Fig. S26). All isoforms were quantified using six different methods. However, significant differences between HD and WT samples were detected only by RSEM and StringTie for the transcript Ndufs4_ENSMUST00000225707.2, and by Salmon for Ndufs4_ENSMUST00000225978.3. The other methods showed no significant results for any isoforms. To explore this discrepancy, we profiled the consecutive 50-mer sequencing counts assigned to each isoform across three HD and three WT samples (as depicted in Fig. 6e and supporting information Fig. S27), alongside the 50-mer global overlapping degree. Regions with a minimal shared degree of 1 represent sequences unique to a specific isoform, such as Ndufs4_ENSMUST00000225707.2. The GaussF method quantifies transcript abundance by focusing solely on unique regions, whereas other methods must account for reads mapped across nearly the entire transcript, which are then probabilistically assigned to overlapping isoforms. This assignment process inherently introduces quantification uncertainty, as it depends on probabilistic models to manage ambiguous reads. For instance, sequencing counts within the unique region of Ndufs4_ENSMUST00000225707.2 are exceedingly low-close to zero-particularly in HD samples (as illustrated in Fig. 6e, insert). In stark contrast, adjacent regions with higher isoform overlap exhibit substantially elevated sequencing counts. Consequently, even minor variations in the probabilistic assignment of ambiguous reads by traditional methods can result in substantial discrepancies in transcript quantification.

The distinct abundance values for the same individual Ndufs4 isoform, observed across different quantification methods, highlight how method-determined abundance estimates can arise from the same raw dataset (as shown in Fig. 6d and Supporting Information Figure S26). Method-specific probabilistic algorithms for read assignment introduce inherent uncertainty, complicating downstream analyses and leading to potential confusion or controversy in interpreting quantification results. Similarly, the neuron death-associated gene Ngfr (ENSMUST00000000122.7) exhibits significant assignment ambiguities (see Fig. 6f). A detailed 50-mer sequencing coverage analysis of Ngfr reveals a sudden spike in the overall profile, corresponding to a region shared by multiple isoforms (as illustrated in Fig. 6g). In this region, the uncertain assignment of ambiguous reads can result in significant inaccuracies in isoform quantification. Compared to the very high sequencing counts induced by multiple-isoform overlaps, the widespread minimal-shared regions, characterized by a global overlapping degree of 1, are associated with much lower sequencing counts. The GaussF method effectively addresses this issue by focusing exclusively on unique regions, thereby eliminating ambiguity. In addition to the genes identified by traditional methods in GO analysis, GaussF-based analysis also identifies genes involved in biological processes, such as protein folding (as illustrated in Fig. 6c). Specifically, the Cct7 gene was identified during biological processes enrichment for HD. Cct7 plays a crucial role in facilitating the folding of specific subsets of proteins.[36] All six detectable Cct7 isoforms were quantified using six different methods (as shown in Fig. 6h), but no consistent quantification pattern was observed across the methods for any isoform. Specifically, Cct7_ENSMUST00000204592.3 was not detected by GaussF or Cufflinks in any of the three HD or three WT samples. Additionally, Cct7_ENSMUST00000204489.3 was not detected by GaussF. However, these isoforms were detected by other methods. This method-induced variability underscores the complexity and uncertainty inherent in transcript quantification, often leading to confusion and controversy. Additionally, GaussF identified two isoforms-Cct7_ENSMUST00000205096.2 and Cct7_ENSMUST00000205143.2-which are enriched in the protein folding process, but other methods didn’t identify such significant changed isoforms. To thoroughly evaluate the quantification of these isoforms, we analyzed the consecutive 50-mer counting profiles and global overlapping degree profiles for all Cct7 isoforms across the three HD and three WT samples. By stacking these profiles (see Fig. 6i and Supporting Information Fig. S28), we aimed to better understand the ground-truth read coverage. The low degree (degree = 1) and low sequencing count within the minimal-shared region contrast sharply with the high degree and high sequencing count within the multiple isoform overlapping regions. This pronounced disparity suggests that the minimal-shared regions provide a more specific and representative measure of the transcript’s abundance. In GaussF, there is no ambiguity in isoform quantification for such regions; however, traditional methods assign all mapped reads to multiple involved isoforms. Moreover, an examination of the 50-mer sequencing coverage for Cct7_ENSMUST00000204592.3 (as shown in Fig. 6j), undetectable by GaussF, reveals that its minimal-shared degree of 1 corresponds to a unique region entirely lacking 50-mer counts across all HD and WT samples. Consequently, GaussF assigns a null quantification to this isoform, and this sharp drop in sequencing coverage supports the conclusion that this isoform is not expressed. In addition, 50-mer sequencing coverage profiles with global 50-mer overlap were generated for the other four Cct7-related isoforms across three HD and three WT samples (as shown in Supporting Information Fig. S29). For these isoforms, the global 50-mer overlap profile generally aligns with the overall sequencing count profile. The sharp drops in sequencing coverage specifically within minimal-shared regions closely correlate with corresponding declines in sequencing counts. These minimal-shared region-specific sequencing count profiles offer valuable insights, suggesting that quantifying transcript abundance based on these regions may yield a more accurate representation of complex transcript abundance. Moreover, GaussF identifies another protein-folding-related gene, Dnajc18, through GO enrichment analysis and quantifies all four of its isoforms. Notably, considerable variation exists among the six methods used to quantify these isoforms (as illustrated in Supporting Information, Fig. S30). This highlights method-induced differences in isoform abundance estimation. For example, all methods except GaussF report a statistically significant difference in the Dnajc18_ENSMUST00000025208 isoform between HD and WT samples. However, the cumulative 50-mer sequencing count across the full length of Dnajc18_ENSMUST00000025208 reveals no significant difference between the three HD and three WT samples (as shown in Supporting Information, Fig. S31). Given that k-mer counts directly correlate with sequencing read counts, this lack of difference in 50-mer counts suggests comparable mapped read counts for this isoform between HD and WT samples. This discrepancy further highlights uncertainties in read assignment during quantification by traditional methods. GaussF’s superior specificity and sensitivity make it more effective in accurately identifying and quantifying isoforms.

## Discussion

In the minimal-sharing evaluation, sudden drops in sequencing counts are associated with sharp decreases in overlapping degree, reflecting a global isoform overlap pattern that influences the corresponding k-mer sequencing count profile. While minimal-shared regions should ideally represent a specific isoform, occasional sharp drops in k-mer counts may arise due to complex sequencing biases. To investigate whether these sharp fluctuations are bias-related, we initially examined the k-mer overlap profile to identify regions with consistent isoform overlap from the same isoform set. This analysis confirms that k-mer counts in these regions originate from a uniform isoform set, so any variations in read coverage reflect sequencing biases alone. Indeed, we observed that complex sequencing biases can cause uneven read or k-mer coverage, though the biased counts tend to vary gradually within regions of uniform isoform-set overlap (see Fig. 6e, 6g, 6i, 6j, S27, S28, S29, and S30). Thus, sharp drops in sequencing counts are more likely due to transcript-specific minimal-shared regions rather than sequencing bias.

Isoform abundance quantification using traditional methods often yields conflicting results, even when applied to the same dataset. For example, in the quantification of Cct7 isoforms (see Fig. 6h), the abundance patterns of Cct7_ENSMUST00000204489.3 and Cct7_ENSMUST00000205096.2 show completely opposite trends across three HD and three WT samples. According to Cufflinks, Cct7_ENSMUST00000204489.3 shows a decrease in abundance from HD to WT, whereas other methods indicate an increase. Such discrepancies can mislead downstream analyses and introduce uncertainty into biological interpretations. Similar contradictions are observed in the quantification of Ndufs4 isoform ENSMUST00000225701.2 (see Fig. S19). These inconsistencies are primarily due to the probabilistic models used in traditional methods, which assign ambiguous reads based on empirical assumptions. In contrast, GaussF’s minimal-shared region approach offers more reliable quantification by reducing ambiguity in read assignment, thereby enhancing the accuracy of isoform abundance estimates and minimizing potential misinterpretations in subsequent analyses.

The minimal-shared region exceeding a degree of 1 indicates a lack of unique sequence for specific transcripts, affecting approximately 30% of human transcript isoforms and 13% of Mus musculus isoforms (see Fig. 4m and 4n). This poses a critical challenge for siRNA knockdown assays targeting these transcripts, as there is no distinct sequence segment to enable specific targeting. When significant changes are observed in such transcripts, careful consideration is necessary for downstream biological assays. For instance, in a knockdown assay, the absence of unique targetable sequences within these transcripts (as illustrated in Supporting Information Fig. S32) means that any chosen siRNA sequence would likely affect multiple isoforms. In terms of quantification, the minimal-shared region-based method offers advantages over traditional probabilistic approaches by simplifying the read or k-mer matching structure. Although these shared regions are common to at least two isoforms, they represent the smallest set of isoforms, thereby improving quantification accuracy. Given the current limitations in accurate count-assignment algorithms, we avoid splitting the quantification value of minimal-shared regions across different transcripts. Thus, the quantification abundance reflects not only the target transcript but also other involved transcripts. This annotation should be explicitly noted at each step in downstream analyses to account for the shared contribution to the quantified abundance. Furthermore, minimal-shared regions facilitate precise targeting with siRNA assays, allowing direct interference with the relevant transcript isoforms. In cases where significant abundance changes involve isoforms with a minimal-shared degree greater than 1, siRNA can effectively target these minimal-shared regions, interfering with all related isoforms simultaneously. The GaussF minimal-shared approach thus seamlessly integrates with downstream experimental assays, enabling exactly targeted interventions.

The k-mer splitting strategy effectively tracks overlap within transcript-specific k-mer sets. When the k-mer length is set to 50 nucleotides, 95.7% of human transcript isoforms share some 50-mer regions with other isoforms, and 64.2% of genes overlap with at least one other gene. Notably, even with longer 500-mer sequences, 72.9% of isoforms and 25.8% of genes still share regions, indicating considerable mapping ambiguity. This analysis uses the Homo_sapiens.GRCh38.cdna.all.fa reference, which includes various transcripts, pseudogenes, and multiple scaffolds, resulting in identical gene names with different gene IDs or gene IDs without direct names. For instance, the genes ENSG00000226165 and ENSG00000232629 both correspond to HLA-DQB2 but have distinct Ensembl Gene IDs because HLA-DQB2 appears on two separate MHC scaffolds due to multiple haplotypes.[37] Ensembl assigns unique identifiers for genes on alternative haplotype scaffolds to account for regions like the MHC, known for high genetic variability. By using an Ensembl reference-based k-mer splitting strategy that retains these alternative scaffolds, the analysis captures the extensive diversity in regions critical for understanding individual genomic differences. This approach highlights gene and isoform overlaps, even amidst the complexities of alternative haplotypes and genomic diversity. Interestingly, as the k-mer length increases from 50 nucleotides to 500 nucleotides, overlap frequency decreases, which helps in distinguishing distinct gene features by reducing background complexity. However, this also reduces the granularity of the analysis, as longer k-mers cover broader regions and may miss smaller, yet biologically significant, transcript segments. This trade-off suggests that choosing an optimal k-mer length involves balancing resolution with specificity: shorter k-mers offer higher resolution for detecting subtle transcript regions, while longer k-mers facilitate feature differentiation at the expense of detail.

In RNA-seq analysis, abundance profiling at the isoform level provides a detailed view of biological function, whereas gene-level profiling offers only a broad, less specific interpretation. For instance, the Homo_sapiens.GRCh38.cdna.all.fa reference contains 30,157 well-annotated genes, compared to 199,385 annotated isoforms, highlighting the greater complexity captured at the isoform level. Importantly, gene-level transcript profiles represent a combined measure of all associated isoforms, which dilutes specificity and reduces the resolution in identifying distinct biological functions. Currently, GO and KEGG pathway analyses primarily rely on gene-level data, treating each gene as a single functional unit. This approach simplifies analysis by assuming that gene-level summaries capture the overall function or pathway involvement. While efficient, this strategy overlooks isoform-specific nuances that may be biologically significant. Moving to isoform-level analyses, although more complex in data processing, would likely enhance the precision and accuracy of functional and pathway insights. This shift is particularly beneficial in cases where isoforms have distinct roles or implications in disease, potentially offering more nuanced interpretations in biological research.

### Conclusions

In summary, our minimal-sharing strategy addresses the long-standing challenge of mapping ambiguity in RNA-seq by isolating transcript-specific k-mers and leveraging minimal-shared regions for precise isoform quantification. Through comprehensive evaluations across multiple species and applications, including an RNA-seq analysis of a Huntington’s disease mouse model, our approach demonstrated superior performance over traditional methods, reducing false positives and negatives while enhancing gene ontology annotations and the detection of key isoforms. By improving the accuracy and reliability of transcriptome quantification, this method provides a robust framework for advancing RNA-seq-based studies of complex biological systems and diseases.

## Methods

### Detailed computational procedure for minimal-sharing strategy

#### 1. Splitting the entire transcript sequence into k-mers

##### 1.1 Generate overlapping k-mers

- Split the entire Exon DNA or mRNA sequence into partially overlapping subsegments (k-mers) of a specified length (e.g., 50 nucleotides), shifting by one base between consecutive k-mers. Example: For an mRNA sequence “ATCGTACG…” with a k-mer size of 50 nucleotides, segment the sequence from positions 1–50, then 2–51, and so on, until the end of the sequence.

##### 1.2 Create an exon or transcript-specific k-mer set

- Compile all generated k-mers from the transcript to form a set for subsequent matching and analysis.

#### 2. Profiling k-mer overlapping degrees

The overlapping degrees of k-mers were evaluated at three levels to provide a comprehensive analysis of their distribution. The global overlapping degree measures the frequency of k-mer occurrences across the entire genomic exon or transcriptome, while the homologous isoform overlapping degree focuses on k-mer occurrences within the same gene family. Finally, the local overlapping degree quantifies the specificity of k-mers to individual isoforms, offering insights into their distinct sequence features.

##### 2.1 Compute the global overlapping degree

- Match each k-mer from the exon or transcript-specific set against all k-mers from all other exon or transcript isoforms in the genome or transcriptome.
- Profile the consecutive k-mers along the exon or transcript based on their global overlapping degrees.

##### 2.2 Compute the homologous isoform overlapping degree

- Match each k-mer against all k-mers from homologous isoforms (isoforms of the same gene or gene family).
- Profile the k-mers along the exon or transcript to reflect their homologous overlapping degrees.

##### 2.3 Compute the local overlapping degree

- Match each k-mer against all k-mers within the same exon or isoform (self-comparison).
- Profile the k-mers along the exon or transcript based on their local overlapping degrees.

#### 3. Implementing the transcript-specific minimal-sharing strategy

##### 3.1 Filter k-mers based on minimal sharing

- From the overlapping degree profiles, identify and retain k-mers with minimal shared degrees.
- Filter out k-mers with higher overlapping degrees to enhance specificity.

##### 3.2 Select minimal-shared k-mer regions

- Recognize that minimal-shared regions may originate from different transcript sets.
- Select the longer minimal-shared region to represent the transcript, ensuring maximum uniqueness and effective representation.

## Data analysis

### 1. Data processing

The data processing pipeline for analyzing raw paired-end sequencing data is meticulously designed to ensure high-quality results through a series of specialized tools tailored for each step of the process. The pipeline begins with an initial quality assessment using FastQC (v0.11.8) from Babraham Bioinformatics, which provides a comprehensive overview of the data’s quality. Following this, adapter sequences are removed, and low-quality bases are trimmed using both Cutadapt (v2.10) and Trimmomatic (v0.39), ensuring that only high-quality data proceeds to the next stage. The trimmed reads are then aligned to the human reference genome GRCh38.p14 (as per Ensembl version) using STAR (v2.7.3a). This dual approach ensures precise and reliable mapping of the reads. Post-alignment, tasks such as sorting and indexing of the mapped reads are efficiently handled by Samtools (v1.7). Additionally, the alignment of k-mer segments to transcript references based on Ensembl annotations enhances the granularity of the genomic insights obtained. Together, these tools form a comprehensive and robust pipeline that ensures meticulous preparation, accurate alignment, and insightful genomic analysis. This integrated approach guarantees that the datasets are processed with precision, leading to reliable and meaningful results.

### 2. Gene level k-mer overlapping analysis

#### 2.1 Exon k-mer overlapping frequency analysis

The *exon_kmer_frequency_distribution.py* script performs a comprehensive analysis of k-mer occurrences at the exon level within genomic data. This Python tool extracts exon sequences, examines overlapping k-mer substrings, and outputs the results in CSV format, streamlining k-mer-based studies on exon structure. Initially, it parses a GTF file to gather details about each exon’s location, strand orientation, and identifiers-crucial elements for accurately pinpointing sequences in the genome. For exons located on the reverse strand, it calculates the reverse complement to maintain strand specificity. Leveraging a FASTA genome file, the script retrieves the exact nucleotide sequence for each exon, then divides it into overlapping k-mer segments. It computes both local k-mer frequencies (unique to each exon) and global frequencies (across all exons) and saves this data in CSV files, each documenting k-mer sequences, their frequencies, and associated transcripts for each exon. This systematic approach supports in-depth analyses of exon structures in genomic studies. The Python script *kmer_frequency_overlapping_exons.py* processes a set of CSV files containing exon and gene information, calculating and updating specific frequency values for each entry. It iterates through CSV files in a specified input directory, extracts exon and gene details, and performs calculations to determine the frequency differences between global and local occurrences. It also counts unique gene occurrences based on the data within each file.

#### 2.2 Transcript k-mer frequency analysis

The script *kmer_frequency_allgenes.py* is designed to analyze k-mer occurrences within a FASTA file of transcript sequences, providing a comprehensive k-mer frequency analysis. It systematically extracts k-mers from each transcript, calculating both their local and global frequencies, and identifies the transcripts in which they appear. For each gene, the script generates CSV files that detail the specific k-mer sequence, its local frequency (the total count of the k-mer across all transcripts of the same gene), its global frequency (the number of distinct transcripts across all genes that contain the k-mer at least once), and a list of all transcripts from all genes that contain the k-mer.

#### 2.3 K-mer overlapping analysis

The script, kmer_frequency_overlapping.py, processes CSV files containing k-mer data from gene-specific outputs to analyze and update frequency information related to gene isoforms. It identifies files using a specific pattern and calculates the homologous gene overlapping frequency (Samegene_Frequency), which measures the presence of k-mers in transcripts of the same gene. Additionally, it computes the heterologous gene overlapping frequency (overlapping_diff_frequency) by finding the difference between the global frequency and the same-gene frequency. The script also identifies the gene_set, encompassing all related gene names involved in the analysis.

#### 2.4 Max heterologous gene frequency

The script Max_Overlapping_Diff_Frequency.py outputs the maximum number of heterologous genes corresponding to each gene’s k-mers.

### 3. Isoform Level k-mer overlapping Analysis

#### 3.1 Isoform k-mer frequency distribution

The script kmer_frequency_distribution.py is designed to analyze and catalog k-mer sequences in transcript sequences from a FASTA file containing human cDNA data. It begins by reading the transcripts and storing them, then processes each transcript to calculate global k-mer frequencies across all transcripts and stores which transcripts contain each k-mer. In the next phase, the script generates a CSV file for each transcript, documenting each k-mer’s local frequency (within that transcript), global frequency (across all transcripts), and listing the specific transcripts where the k-mer is found. This allows for detailed comparison of k-mer presence and frequency across isoforms, aiding in genomic analysis and comparative studies.

#### 3.2 Comprehensive isoform k-mer frequency analysis

kmer_frequency_overlapping_isoforms.py processes a set of CSV files located in the specified directory to identify the maximum value of the overlapping_diff_frequency column in each file. It begins by searching for all CSV files within the target directory, then iterates through each file to read its contents and calculate the maximum value in the gene_set column (assumed to be a representation of overlapping differences). These maximum values are stored in a list, along with the respective file names.

#### 3.3 Max heterologous gene frequency for isoforms

Max_Overlapping_Diff_Frequency_isoform.py outputs the maximum number of heterologous isoforms corresponding to each isoform.

#### 3.4 The minimal-shared k-mer set filtering

kmer_frequency_distribution_mini_shared.py script processes a FASTA file containing nucleotide sequences, breaking each entire transcript isoform sequence down into k-mers (subsequences of a specified length). It calculates both local frequency (how often each k-mer appears within the given transcript isoform) and global frequency (how often each k-mer appears across all other transcript isoforms) for these k-mers. The script identifies k-mers with the minimum global frequency for each transcript, tracks which transcripts contain each k-mer, and writes this information into individual CSV files for each transcript. The script employs command-line arguments to specify input and output paths and uses tqdm for progress tracking.

#### 3.5 k-mer counting from sequencing data

This Python script kmer_count_v2_20240803.py is designed for processing large FASTQ files, specifically for counting occurrences of k-mers (subsequences of length k) in sequencing data and matching those counts against a list of k-mers from CSV files. It operates by leveraging multithreading for efficient parallel processing, where a producer function reads the FASTQ file in chunks and extracts sequences, while multiple consumer threads process these sequences into k-mers and count their frequencies. The script uses a Queue to manage communication between the producer and consumers, ensuring memory efficiency. After counting k-mers, it matches the counts with a set of specified k-mers from input CSV files, generating output CSVs that contain the frequency of each k-mer, making it suitable for large-scale genomic data analysis.

#### 3.6 Merging and normalizing k-mer counting data

This Python script merge_normalizing_count_TPM.py is designed to merge k-mer count data with a reference k-mer dataset and calculate normalized k-mer counts for each entry. Given a directory of k-mer reference CSV files and a corresponding directory of k-mer count files, it processes each pair of files by merging them on the k-mer column, calculates a normalized frequency for each k-mer based on a specified read length and k-mer size, and saves the resulting merged data into an output directory. The script uses command-line arguments to specify input directories, output directory, read length, and k-mer length, making it adaptable for various data sets and sequencing configurations.

#### 3.7 The transcript isoform quantification

This Python script GaussF_TPM.py is designed to analyze GC content in k-mer data across multiple gene transcript files and fit a Gaussian cumulative distribution function (CDF) to model the data. It processes multiple CSV files in a specified input directory, where each file contains k-mer data for different genes or transcripts. The script calculates the GC content for each k-mer, then fits a Gaussian CDF to various cumulative distributions (such as local frequency, normalized count, and absolute count) within the data. It extracts essential information like gene names and transcript IDs from filenames, applies fitting criteria to ensure meaningful results, and summarizes each file’s fitting parameters in an output CSV file. This streamlined approach enables comprehensive GC content analysis with robust error handling and suitability checks to ensure the reliability of Gaussian fitting for normalized k-mer data.

#### 3.8 Integrated pipeline for merging steps 3.5-3.7 to output isoform tpm quantification

This script pipeline_isoform_quantification.sh automates the process of k-mer counting, merging, and quantification for isoform-specific analysis on a set of trimmed FASTQ files specified in a sample list. The script performs three main tasks: (1) counts isoform-specific k-mers in each sample’s FASTQ file using a specified k-mer counting script, (2) merges the k-mer counts with reference data for normalization, and (3) applies Gaussian filtering for GC-content-based isoform quantification. It supports parallel processing with a defined maximum of concurrent jobs, enhancing efficiency for large datasets. Intermediate files are managed carefully, with directories created as needed and temporary data optionally removed after processing. This setup provides a streamlined approach for high-throughput isoform quantification.

### 4. K-mer chromosomal positioning and gene identification

This script chromosome_position.py is designed to identify specific DNA sequences within a FASTA file and map them to corresponding genes using a GTF file. It first scans through a given FASTA file (Homo_sapiens.GRCh38.dna.primary_assembly.fa) to locate all occurrences of a specified DNA sequence. For each match, the script extracts the chromosome, start, and end positions of the sequence. It then cross-references these positions with gene annotations in the GTF file (Homo_sapiens.GRCh38.112.gtf). This process helps in associating specific DNA sequences with potential genes, aiding in genomic research and analysis.

### 5. Venn diagram and upset plot

The R script venn_upset_method.R script aims to analyze and visually compare gene expression results from multiple RNA-Seq quantification methods, including GaussF, Salmon, Kallisto, RSEM, Cufflinks, and StringTie. After loading the data, the script extracts unique sets of gene identifiers for each method, excluding any missing values, and organizes these sets into a list. To visualize intersections across more than five sets, it employs two approaches: a Venn-like diagram using ggVennDiagram for a simple overview and an UpSet plot using UpSetR for detailed intersection insights, which is particularly effective when handling numerous sets.

### 6. Differential expression analysis

DEseq2_isoform_GaussF.R is designed for differential gene expression analysis using DESeq2. The script begins by loading raw count data from a CSV file, which contains gene expression values across multiple replicates for each group. It preprocesses the data by standardizing transcript identifiers, ensuring unique rows, and filtering low-expression genes to focus on those expressed in a significant number of samples. Then, it creates a count matrix and sample information. After setting up the DESeq2 dataset, the script filters out low-count genes, runs the DESeq function to obtain differential expression results, and removes any rows with NA values. Finally, it adds gene identifiers back to the results and saves the output to a new CSV file.

### 7. GO and KEGG analysis

GO_KEGG_threshold_individual.R performs functional enrichment analysis on differentially expressed genes (DEGs) obtained from RNA-seq data. The script begins by reading the DEG data from a CSV file, verifying the presence of key columns (gene ID, log2 fold change, and p-value), and filtering genes based on significance thresholds (p-value < 0.05 and |log2FoldChange| > 1). The filtered gene list is converted to ENTREZ IDs, essential for downstream analysis. The script then conducts Gene Ontology (GO) enrichment analysis across three categories-Biological Process (BP), Molecular Function (MF), and Cellular Component (CC)-and generates visualizations (bar plots and dot plots) for enriched terms in each category. Additionally, the script performs KEGG pathway enrichment analysis, visualizing significant pathways and saving all results into dynamically named CSV files for easy reference.

### 8. Data combination and filtering

This Python script GaussF_data_combined.py is designed to consolidate k-mer count data from multiple CSV files located in a specified directory. It sequentially reads each CSV file matching the pattern results_file_GC_*.csv, extracts relevant columns (such as File, Gene_Name, Transcript_ID, Global_Frequency, Present_in_Transcripts, Transcript_Length, and Report), and retrieves the “Sum or Fitted A (Abundance) for Normalized Count” column, renaming it based on the file name for clarity. The script then merges each processed file into a single DataFrame, ensuring alignment on common columns, and outputs the final combined dataset to a new CSV file. Scripts like stringtie_data_combined.py, salmon_data_combined.py, kallisto_data_combined.py, RSEM_data_combined.py, and cufflinks_data_combined.py are used for combining data from various sources.

### 9. Data filtering

GaussF_data_filter.sh script is designed to extract specific rows from a CSV file, based on transcript IDs listed in a text file. It begins by emptying the designated output file to ensure a clean start. The script then reads each transcript ID from the input list and searches for matches in the third column of the CSV file using awk. Each matching row is appended to the output file.

### 10. Minimal shared region extraction

output_summary.py extracts the global frequency of transcripts and isoforms in the minimal shared region and the corresponding isoforms.

### 11. Pie chart and histogram plotting

pie_chart_plot.py creates pie charts and histograms for isoforms corresponding to the minimal shared region, indicating unique, shared between two isoforms, or shared among three isoforms.

### 12. ID conversion

ENTREZID_to_ensembleid.R converts ENTREZ IDs obtained from GO and KEGG analysis to Ensemble IDs.

### 13. PCA plotting

PCA_isoform_GaussF.R script is designed to process and analyze gene expression count data for differential expression and principal component analysis (PCA) using DESeq2. The target data is a CSV file containing gene counts from samples. The script first preprocesses the data by cleaning and standardizing the “Present_in_Transcripts” column, ensuring unique entries and selecting relevant columns for analysis. It then filters genes based on mean expression levels, removes any rows with missing values, and converts the data into a count matrix format suitable for DESeq2. Sample information is used to create a DESeq2 dataset. After performing variance stabilizing transformation (VST), the script generates a PCA plot with ggplot2, visually separating the two groups. Finally, PCA coordinates are saved to a text file for further analysis or documentation. Scripts like PCA_isoform_GaussF.R, PCA_isoform_RSEM.R, PCA_isoform_Cufflinks.R, PCA_isoform_Salmon.R, PCA_isoform_Kallisto.R, and PCA_isoform_Stringtie.R are used for PCA plotting of isoforms from different data sources.

### 14. Generating rna-seq simulation data using polyester

This R script polyester_simulated_data.R is designed for simulating RNA-Seq data by using a reference transcriptome, abundance data, and specific simulation parameters. It installs required packages, loads the reference transcriptome file, extracts transcript IDs, and aligns them with abundance data to create an abundance matrix. The script then sets up a simulation environment, including fold changes, output directory, and checks for essential dependencies.

### 15. Transcript quantification with GaussF

Step-by-step procedural description for isoform aligning by GaussF:

1. Isoform Splitting Based on k-mer Setting (e.g., 50 nt):

- Step 1.1: Each isoform sequence is divided into 50-nucleotide segments, with each segment overlapping the previous one by 49 nucleotides, resulting in a single-nucleotide shift between adjacent segments.

- Example: Given an isoform sequence (e.g., “ATCGTACG…”) and a specified k-mer size of 50 nucleotides, the sequence is segmented into overlapping 50-nucleotide segments, or k-mers. This segmentation starts from the first nucleotide position, creating the initial k-mer spanning positions 1-50. For each subsequent k-mer, the window shifts by one nucleotide, producing segments that span positions 2-51, 3-52, and so on, until reaching the end of the sequence. This approach generates a series of overlapping k-mers, each offset by a single base, that collectively cover the entire sequence.
- Step 1.2: Ensure that every isoform is processed uniformly to generate a complete set of 50-mers for each. This consistent approach ensures comprehensive coverage across all isoforms, with each sequence segmented into overlapping 50-nucleotide k-mers.
2. Whole Transcriptome-Level Isoform 50-mer Alignment Process:

- Step 2.1: Gather all 50-mer sequences generated in step 1 from every isoform across the entire transcriptome.
- Step 2.2: Perform an alignment of these 50-mers against the full set of isoform 50-mers to identify matching sequences. This process includes:

`o` Step 2.2.1: Align 50-mers within the homologous gene’s isoforms to identify regions of similarity and distinctiveness.
`o` Step 2.2.2: Matching 50-mer sequences from isoforms of heterologous genes to explore potential cross-gene similarities or shared motifs.
3. Transcript-Specific Minimal Shared 50-mer Regions:

- Step 3.1: Analyze the alignment results to identify the minimal set of shared k-mers that are present across all isoforms.
- Step 3.2: Use these minimal shared 50-mers to represent the transcript for downstream quantification from sequencing data.

Step-by-Step Procedural Description for Self-benchmarking Gaussian Fitting:

1. Initial Data Preparation:

- Step 1.1: Generate an initial table for each 50-mer along the minimal shared transcript region. This table should include:

`o` Step 1.1.1: 50-mer Sequence Identification. For each 50-mer within the minimal shared transcript region, extract and record the unique 50 nucleotide sequence. This sequence serves as the basis for subsequent analyses and calculations.
`o` Step 1.1.2: Modeling Count Assignment. Initially set the modeling count to 1 for each 50-mer sequence. This provides a standardized baseline.
`o` Step 1.1.3: 50-mer Count from Clean Sequencing Data. Retrieve the sequencing count for each 50-mer based on processed (cleaned) sequencing data. This value represents the observed read frequency for each sequence in a given experimental dataset.
`o` Step 1.1.4: Normalized Sequencing Count (RPKM or TPM). Normalize each 50-mer’s sequencing count to either RPKM (Reads Per Kilobase per Million mapped reads) or TPM (Transcripts Per Million). This step adjusts counts for sequence length and sequencing depth, enabling cross-sample comparison. For RPKM, the formula adjusts for transcript length and total mapped reads, while TPM scales to overall transcript abundance.
`o` Step 1.1.5: GC Content Calculation. Calculate the GC content for each 50-mer by determining the proportion of G and C nucleotides within the sequence.
2. Example Dataset:

- Step 2.1: Demonstrate the method for analyzing 50-mers with a simplified dataset through the process using a simplified dataset consisting of 10 50-mers.
3. Modeling Data Grouping:

- Step 3.1: Group the 50-mers by their GC content, organizing sequences with identical GC content values together.
4. GC Content Distribution:

- Step 4.1: Construct a Gaussian distribution using the GC content groupings, aligning it with the natural distribution pattern of GC content. This approach allows the GC content data to reflect its inherent variability and typical distribution.
5. Gaussian CDF Distribution:

- Step 5.1: The Gaussian Cumulative Distribution Function (CDF) is derived from the Gaussian (or normal) probability density function (PDF).
6. Parameter Estimation:

- Step 6.1: Identify the key parameters-mean and standard deviation-of the GC content distribution using the Gaussian cumulative distribution function (CDF). These parameters are fundamental to characterizing the GC content distribution and are intrinsic, independent of any empirical data.

After processing the sequencing data, the counts are grouped according to the GC content of each 50-mer. These grouped counts are then plotted against the GC content in a cumulative distribution function (CDF) format. Due to various complex biases, sequencing counts for each 50-mer can vary significantly. To address this, a Gaussian CDF with fixed parameters is applied to fit the count distribution across the GC content range. In this fit, the amplitude (A) of the Gaussian represents an abundance estimate, which helps ensure that the assigned counts-or RPKM/TPM values-of individual 50-mers contribute accurately to the final abundance estimation.

1. Grouping and Plotting:

- Step 7.1: Organize the sequencing counts based on the GC content of each 50-mer.
- Step 7.2: Plot the sequencing count against the GC content and in CDF format to visualize the distribution.
2. Fitting Gaussian Distribution:

- Step 8.1: Apply a Gaussian cumulative distribution function (CDF) with fixed parameters to fit the sequencing count plot as a function of GC content. In this fitting, the amplitude *A* represents the abundance estimate.
3. Abundance Estimation:

- Step 9.1: Use the amplitude (A) obtained from the Gaussian fit to represent the abundance.

### Collection of tissue samples

Human colorectal samples were ethically obtained from the Sun Yat-sen University Cancer Center and Shenzhen University General Hospital. Before sample collection, all donors provided informed consent, permitting the acquisition of biopsies and the comprehensive molecular profiling of their transcriptomes.

The B6-hHTTCAG130 mice (B6/JGpt-Tg(hHTT-CAG130)90/Gpt), which carry a human HTT gene fragment with 130 CAG repeat expansions, were obtained from GemPharmatech (Strain No. T054804). All mice were housed in a pathogen-free facility, provided with autoclaved food, and maintained on acidified, autoclaved water. All animal procedures adhered to the guidelines set by the Animal Care and Use Committee of Shenzhen University of Advanced Technology.

### Library preparation

The RNA-seq library preparation methods for both mouse and colorectal samples utilize a common protocol based on the VAHTS Universal V8 RNA-seq Library Prep. This protocol includes several critical steps: RNA fragmentation, hexamer-directed cDNA synthesis, end repair, dA-tailing of fragments, adaptor ligation, PCR library amplification, and subsequent sequencing. Notably, when working with spike-in samples, an alternative tagmentation step is introduced, replacing the end repair, dA-tailing, and adaptor ligation steps.

### RNA isolation

Total RNA was extracted from cells using the RNAiso Plus kit (catalog no. 9109, TaKaRa Biotechnology) according to the manufacturer’s instructions. The extracted RNA was resuspended in RNase-free water, which was used consistently in all RNA-related assays. To evaluate RNA quality, a 2100 Bioanalyzer with an RNA Pico chip was utilized. The total RNA was then aliquoted into 5 μg portions and stored at -80°C for long-term preservation, ensuring its availability for future experiments.

### rRNA depletion

To accurately profile non-rRNA molecules in RNA samples, we utilized the Ribo-off rRNA Depletion Kit (Human/Mouse/Rat) from Vazyme (kit no. N406-01). This kit provides a reliable and efficient method for removing ribosomal RNA (rRNA) from the total RNA pool. The rRNA depletion process involves several key steps: First, the total RNA sample is combined with rRNA removal probes specifically designed to target and bind to rRNA molecules. The mixture is then incubated, allowing the probes to effectively hybridize with the rRNA. After incubation, a removal solution is added to degrade the rRNA-probe complexes. The remaining RNA, now enriched with non-rRNA molecules, is purified to obtain the desired RNA pool for downstream applications. By using the Ribo-off rRNA Depletion Kit, we significantly reduce the abundance of rRNA, enabling a more comprehensive analysis of the non-rRNA fraction and enhancing the sensitivity of subsequent RNA-seq experiments. This rRNA depletion step is crucial for obtaining high-quality data, ultimately contributing to the accuracy and reliability of our molecular analyses.

### Reverse transcription

The rRNA-free RNA samples underwent reverse transcription using random hexamers and SuperScript™ IV reverse transcriptase (Invitrogen, Cat. No. 18090200), in accordance with the manufacturer’s guidelines. The reaction was set up in a 20 μL system, comprising 100 ng of RNA template, 2.5 μM random hexamers, a dNTP Mix (10 mM each), and 1 μL (200 units) of SuperScript™ IV. The mixture was initially incubated at 25°C for 10 minutes, followed by 42°C for 50 minutes to facilitate cDNA synthesis. To terminate the reaction, the mixture was heated to 70°C for 15 minutes, effectively inactivating the reverse transcriptase.

### PCR amplication

The library was amplified using PCR with the HIFI KAPA master mix (2×) in the following composition: 25 μL of 2× HIFI KAPA master mix, 10 μL of cDNA, 13 μL of H2O, and 1 μL of either the universal forward or reverse primer (10 μM concentration).

Universal Forward Primer: 5’-AATGATACGGCGACCACCGAGATCTACACCTCTCTATACACTCTT-3’ (:phosphorothioate)

Universal Reverse Primer: 5’-CAAGCAGAAGACGGCATACGAGATGTGACTGGAGTT-3’ (:phosphorothioate)

The PCR reaction was conducted in a thermocycler with the following conditions: an initial denaturation at 95°C for 5 minutes, followed by 10-15 cycles of 95°C for 15 seconds and 60°C for 30 seconds. The amplified library was then purified using 1.8× Ampure XP DNA Beads and resuspended in 20 μL of H2O. For quality assessment, 1 μL of the library was analyzed using the Qubit 4 Fluorometer with the dsDNA HS (High Sensitivity) Assay Kit from Invitrogen.

### Sequencing

The purified PCR libraries were sequenced using either the Illumina NovaSeq 6000 (PE150) platform or the MGISEQ-2000 (PE150) platform. The data output was generated according to the predetermined yield specifications.

### Statistics

To assess potential deviations between the different samples, we employed non-parametric tests, specifically the Kruskal-Wallis test. The Kruskal-Wallis test is particularly effective for comparing samples with limited size. This approach provides a comprehensive understanding of the relationship between the variables by focusing on their rank order rather than their specific values.

## Author contributions

Qiang Su conceived the study and performed the data analyses, as well as generated the figures. Qiang Su and Yi Long wrote the manuscript, incorporating input from all authors. Deming Gou, Qizhou Lian, Fuyu Duan, Shaoxiang Chen, Hongxiang Zeng, Huanhuan Joyce Chen, and Junmin Quan contributed to the analysis discussion. All authors have read and approved the final version of the manuscript.

## Acknowledgement

The authors thank all members of the Lian lab for helpful discussions.

## Supplementary data

Figs. S1 to S32

## Data availability

The sequencing raw data and processed data have been deposited in the NCBI Sequence Read Archive (https://www.ncbi.nlm.nih.gov/sra) under the accession number PRJNA999048. Specifically, the SRA accession numbers for the RNA-seq data based on S1, S2, and S3 are SRR25438891, SRR25438890, and SRR25438889, respectively. The RNA-seq data for B6-hHTTCAG130 (HD) and B6 (WT) mice, comprising three HD and three WT samples, is available in the Sequence Read Archive (SRA) under project accession PRJNA1185396. The specific SRA accession numbers for the HD samples are musHD_STR-1 (SRR31321647), musHD_STR-2 (SRR31321646), and musHD_STR-3 (SRR31321645), while the WT samples are labeled musWT_STR-1 (SRR31321644), musWT_STR-2 (SRR31321643), and musWT_STR-3 (SRR31321642), enabling detailed comparative analysis between HD and WT mouse models.

## Code availability

The code used for data analysis is available at https://github.com/QiangSu/Minimal-Sharing-Strategy.

